# Human peroxiredoxin 6 is essential for malaria parasites and provides a host-based drug target

**DOI:** 10.1101/2022.04.11.487889

**Authors:** Matthias Paulus Wagner, Pauline Formaglio, Olivier Gorgette, Jerzy Michal Dziekan, Christèle Huon, Isabell Berneburg, Stefan Rahlfs, Jean-Christophe Barale, Sheldon I. Feinstein, Aron B. Fisher, Didier Ménard, Zbynek Bozdech, Rogerio Amino, Lhousseine Touqui, Chetan E. Chitnis

## Abstract

The uptake and digestion of host hemoglobin by malaria parasites during blood stage growth leads to significant oxidative damage of membrane lipids. Repair of lipid peroxidation damage is crucial for parasite survival. Here, we demonstrate that *Plasmodium falciparum* imports a host antioxidant enzyme, peroxiredoxin 6 (PRDX6), during hemoglobin uptake from the red blood cell cytosol. PRDX6 is a lipid peroxidation repair enzyme with phospholipase A_2_ (PLA_2_) activity. Inhibition of PRDX6 with a PLA_2_ inhibitor, Darapladib, increases lipid peroxidation damage in the parasite and disrupts transport of hemoglobin-containing vesicles to the food vacuole, causing parasite death. Furthermore, inhibition of PRDX6 synergistically reduces the survival of artemisinin-resistant parasites following co-treatment of parasite cultures with artemisinin and Darapladib. Thus, PRDX6 is a unique host-derived drug target for development of antimalarial drugs that could help overcome artemisinin resistance.

**GRAPHICAL ABSTRACT:** 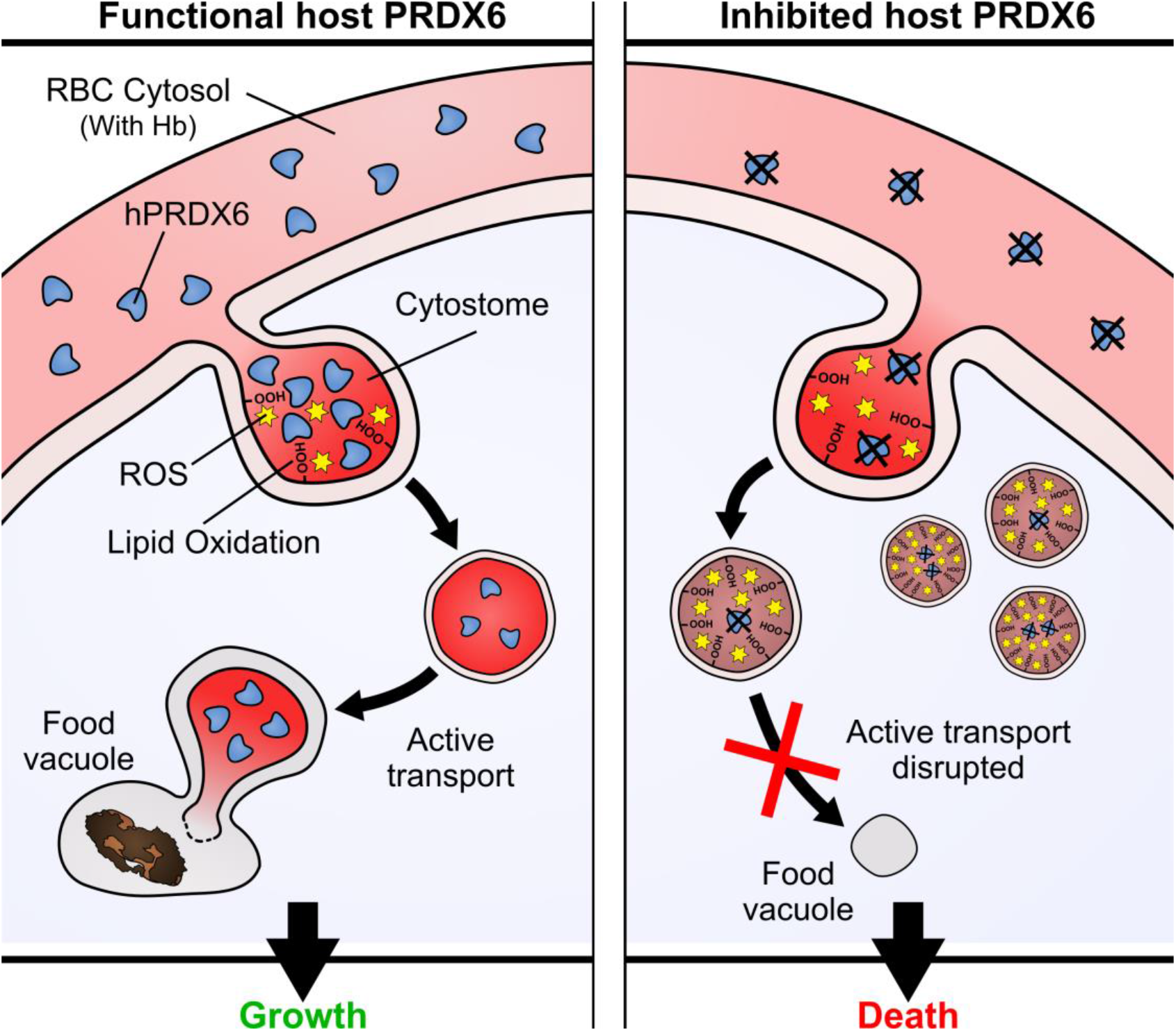

## INTRODUCTION

The mosquito-borne apicomplexan parasite, *Plasmodium falciparum*, was responsible for about 229 million malaria cases in 2019 that led to an estimated 409 000 deaths (WHO, 2020). Malaria thus remains a major public health problem in the tropical world. All clinical symptoms of malaria are attributed to the blood stage of the parasite’s life cycle where *P. falciparum* invades and multiplies within host red blood cells (RBCs). Resistance to the current state-of-the-art antimalarial drug artemisinin (ART) in *P. falciparum*, which was first reported in Southeast Asia in the early 2000s (Noedl et al., 2008) and has now been detected in sub-Saharan Africa (Balikagala et al., 2021; Uwimana et al., 2021), poses a major threat to malaria control and elimination efforts. Mechanistically, *P. falciparum* attains artemisinin resistance by reducing the uptake of host red blood cell hemoglobin, which in turn leads to lower oxidative stress levels and prevents activation of artemisinin, enabling parasite survival (Birnbaum et al., 2020). Thus, understanding the mechanisms for hemoglobin uptake and oxidative stress management by the parasite can lead to the development of new drugs and intervention strategies that overcome artemisinin resistance and enable progress towards malaria elimination.

During blood stage growth, *P. falciparum* internalizes hemoglobin in an endocytic process known as host cell cytosol uptake (HCCU) (Aikawa et al., 1966; Spielmann et al., 2020). Endocytosed hemoglobin is transported in vesicles to an acidic, lysosome-like parasite organelle termed food vacuole (FV), where hemoglobin is proteolytically digested to provide amino acids for synthesis of parasite proteins (Bakar et al., 2010; Jonscher et al., 2019; Milani et al., 2015). The digestion of hemoglobin releases free heme which is partly detoxified by crystallization into hemozoin (Hempelmann, 2007). The Fe(II) ion within the free heme catalyses the production of reactive oxygen species (ROS) that cause the oxidation of unsaturated membrane phospholipids (PLs) (Bochkov et al., 2010; Das and Nanda, 1999; Fu et al., 2009). This process, termed lipid peroxidation, is toxic for cells as it causes drastic changes to membrane fluidity, rigidity, curvature and permeability (Agmon et al., 2018; Borst et al., 2000; Runas and Malmstadt, 2015). Efficient repair of lipid peroxidation damage is thus crucial for cell survival (Bochkov et al., 2010).

So far, no parasite enzyme that is involved in repair of lipid peroxidation damage during blood stage growth has been identified. However, RBCs contain a highly abundant lipid peroxidation repair enzyme called peroxiredoxin 6 (PRDX6) (Gautier et al., 2018; Li et al., 2015). Mammalian PRDX6 is a conserved trifunctional enzyme with phospholipase A_2_ (PLA_2_), lysophosphatidylcholine acyl transferase (LPCAT) and cysteine-dependent peroxidase activities (Chowhan et al., 2020; Fisher, 2018; Fisher et al., 2016; Li et al., 2015; Wu et al., 2009). The PLA_2_ function of PRDX6, which selectively cleaves oxidized fatty acid side chains and allows subsequent replacement with an unoxidised fatty acid through its LPCAT activity, is crucial for lipid peroxidation damage repair (Bochkov et al., 2010; Fisher, 2018; Fisher et al., 2016).

Here, we show that *P. falciparum* internalizes host RBC PRDX6 along with hemoglobin during blood stage parasite development. We demonstrate that lipid peroxidation repair by PRDX6 is essential for parasite development and growth, identifying PRDX6 as a novel host-derived drug target against malaria parasites. Treatment of *P. falciparum* blood stage cultures with a PLA_2_ inhibitor, Darapladib, inhibits the PLA_2_ activity of PRDX6 and blocks parasite growth. Moreover, co-treatment with Darapladib and artemisinin synergistically reduces survival of ART-resistant *P. falciparum* strains in a ring stage survival assay. The advantage of antimalarial drugs that target host enzymes has been recently highlighted (Wei et al., 2021). The studies reported here identify PRDX6 as a potential host-based drug target and may provide a strategy to overcome artemisinin resistance in *P. falciparum*.

## RESULTS

### *P. falciparum* imports host RBC PRDX6 along with hemoglobin

Given the abundance of PRDX6 in the RBC cytosol, we hypothesized that *P. falciparum* may internalize host PRDX6 along with hemoglobin during host cell cytosol uptake (HCCU). To assess this, we selectively lysed the RBC membrane of *P. falciparum*-infected RBCs by saponin treatment and detected PRDX6 in purified parasites. Saponin-treated *P. falciparum* blood stages were analysed by mass spectrometry to identify internalized host RBC proteins. PRDX6 was one of 97 host RBC proteins that were repeatedly detected in saponin-treated parasite fractions in each of 15 independent experiments (Fig. S1 and Table S1). The presence of PRDX6 in saponin-treated parasite fractions was also examined by western blotting using an anti-PRDX6 mouse monoclonal antibody. PRDX6 was detected in lysates of both intact *P. falciparum* infected RBCs as well as saponin-treated parasite fractions (Fig. S2). At the same time, hemoglobin was detected only in intact parasite-infected RBCs and not in saponin-treated parasite fractions confirming complete lysis of the RBC membrane and loss of the RBC cytosolic fraction (Fig. S2). Localization of human PRDX6 within the parasite in *P. falciparum*-infected RBCs was also confirmed by immuno-fluorescence microscopy. PRDX6 was evenly distributed in the cytosol of uninfected RBCs (Fig. 1A). In contrast, multiple PRDX6-containing vesicles were detected adjacent to the parasite FV in *P. falciparum* infected RBCs suggesting endocytic uptake of PRDX6 by the parasite (Fig. 1A). The subcellular localization of internalized PRDX6 was further confirmed by immuno-electron microscopy (EM). As expected, uninfected RBCs showed uniform cytosolic staining of PRDX6 (Fig. 1B), whereas in infected RBCs, PRDX6 was localized in electron-dense, hemoglobin containing vesicular structures (Fig. 1C-F). PRDX6-containing vesicles were found at the inner surface of the parasite plasma membrane in structures that resembled cytostomes (Fig. 1C-E black arrows), adjacent to the FV (Fig. 1C-E white arrows) and in the parasite cytosol (Fig. 1F striped grey arrow). Taken together, these images demonstrate that host PRDX6 is internalized by *P. falciparum*. Furthermore, the data suggests that PRDX6 is internalized by endocytosis during HCCU and subsequently transported in hemoglobin-containing vesicles (HCV) to the FV. PRDX6 was not detected inside the FV lumen, suggesting that it may be cleaved after delivery to the FV, which contains proteases involved in hemoglobin degradation (Francis et al., 1997).

**Figure 1.**
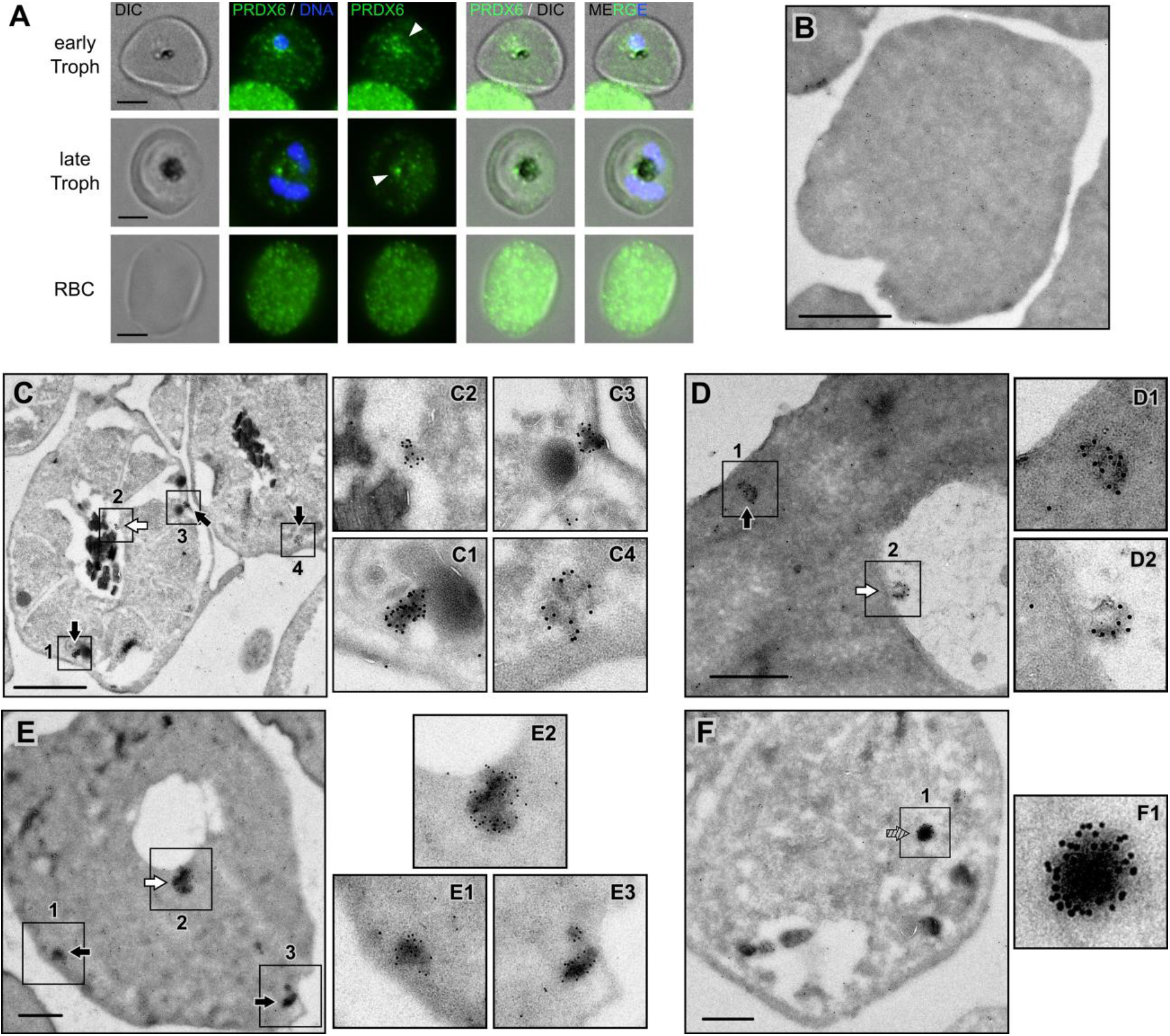
Human PRDX6 is internalized by *P. falciparum* along with hemoglobin during HCCU. (**A**) Immuno-fluorescence microscopy shows even cytosolic staining of PRDX6 in uninfected RBCs. In early and late trophozoites, multiple PRDX6 foci (white arrow heads) were observed adjacent to the parasite FV. DNA, Hoechst 33342 (nuclei). Scale bars: 5 μm. (**B-F**) Immuno-transmission electron microscopy (TEM) using monoclonal mouse anti-PRDX6 antibody. (**B**) Uninfected RBC showing even cytosolic staining for PRDX6. (**C-F**) *P. falciparum*-infected RBCs show co-localization of PRDX6 and hemoglobin in vesicles at the inner surface of the parasite plasma membrane in structures that resemble cytostomes (black arrows), at the FV (white arrows), or within the parasite cytosol (striped grey arrow). Scale bars: 500 nm. Representative images from three independent experiments are shown.

### The PLA_2_ inhibitor Darapladib inhibits PRDX6 and blocks blood stage growth by impairment of lipid peroxidation repair

To investigate the role of PRDX6 during blood stage growth, inhibitors of structurally and functionally related PLA_2_ enzymes were screened for binding to the PLA_2_ active site of PRDX6 by activity-based protein profiling (ABPP) using recombinant human PRDX6 (Fig. S3) and the fluorescently tagged probe TAMRA-FP that covalently labels the serine in the PLA_2_ active site of PRDX6 (Fig. 2A, Fig. S4) (Cravatt et al., 2008). Inhibitors that bind the PLA_2_ active site of PRDX6 compete with TAMRA-FP and reduce the fluorescent labelling of PRDX6. A range of PLA_2_ inhibitors were tested for binding using ABPP (Fig. 2A, Fig. S4). While the inhibitors ATK, MAFP and Darapladib bound PRDX6, Varespladib and P11 failed to bind. The different PLA_2_ inhibitors were also tested *in vitro* for their ability to block *P. falciparum* blood stage growth and progression from rings to schizonts. Synchronous *P. falciparum* 3D7 cultures were treated at the ring stage and effects on ring to schizont development (“progression”) and multiplication following completion of a full blood stage cycle (“growth”) were assessed by flow cytometry using the DNA intercalating fluorescent dye SYBR Green I (Fig. 2B). Notably, inhibitors that bound PRDX6 (ATK, MAFP and Darapladib) blocked parasite progression and growth whereas inhibitors that did not bind PRDX6 (P11 and Varespladib) had no effect. All effective inhibitors arrested parasite development at the trophozoite stage (Fig. 2C). Darapladib had the lowest IC_50_ for parasite progression (0.56 μM) and growth (0.76 μM) (Fig. 2B and Table S1) and was selected for further studies. As expected, Darapladib inhibited the PLA_2_ activity of PRDX6 in a PLA_2_ activity assay (Fig. 2D).

**Figure 2.**
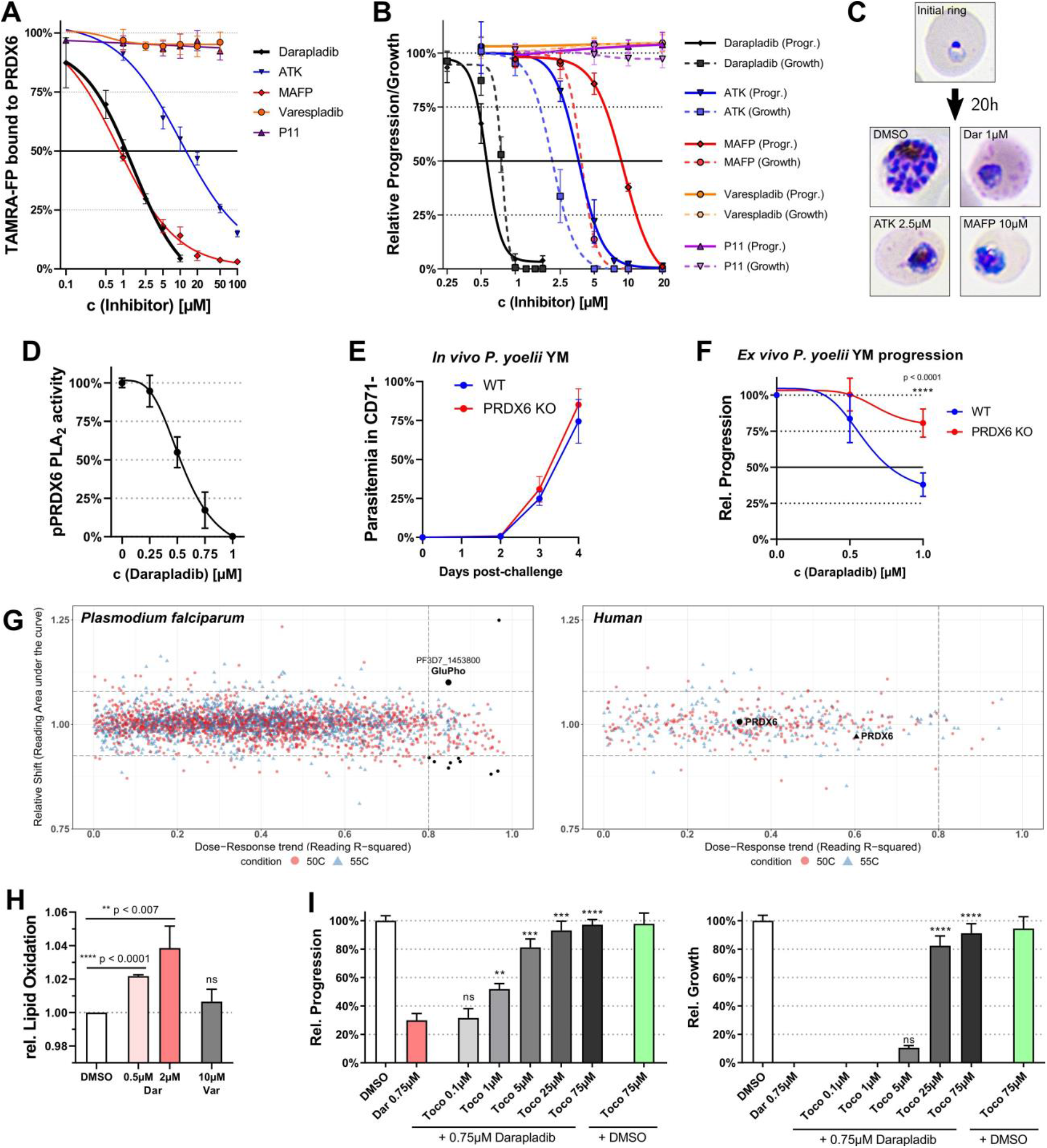
Darapladib selectively inhibits PRDX6 and blocks *P. falciparum* growth by impeding lipid peroxidation repair. (**A**) Binding of TAMRA-FP with recombinant PRDX6 in the presence of different PLA_2_ inhibitors in activity-based protein profiling (ABPP) assays. Reduced labelling of PRDX6 by TAMRA-FP identifies PLA_2_ inhibitors that bind PRDX6, cf. Fig. S4. (**B**) Inhibition of *P. falciparum* ring to schizont progression and blood stage growth with PLA_2_ inhibitors. Parasites were treated at ring stage and growth to schizont stage in the same cycle (“progression”) or to next generation rings (“growth”) was scored by flow cytometry using DNA intercalating fluorescent dye SYBR Green I. (**C**) Representative light microscopy images of control and treated *P. falciparum* blood stages showing arrest at the trophozoite stage following treatment with different inhibitors at ring stage. (**D**) Radioactive PLA_2_ activity assay using phosphorylated recombinant human PRDX6 in presence or absence of Darapladib. Darapladib inhibits PLA_2_ activity of PRDX6 with IC_50_ of ≈0.5μM (**E**) Flow cytometric measurement of *in vivo* growth curves of *P. yoelii* YM in C57BL/6 WT and *prdx6^-/-^* mice. *In vivo* infection was performed two times independently with five mice per group. (**F**) *Ex vivo* ring to schizont progression of *P. yoelii* YM in CD71^-^ mature RBCs from WT and *prdx6^-/-^* mice in presence and absence of Darapladib. Darapladib inhibits *P. yoelii* YM progression from ring to schizont stage in WT RBCs at concentration of 1 μM but not in *prdx6^-/-^* mouse RBCs. (**G**) ITDR-CETSA analysis of protein target engagement by Darapladib (0-100 μM) in trophozoite lysate. Distribution of protein stabilizations, under 50°C (red circle) and 55°C (blue triangle) thermal challenges, is plotted as a function of R^2^ value (goodness of curve fit) against ΔAUC (area under the curve of heat-challenged sample normalized against non-denaturing 37°C control) for all proteins detected in the assay. Three times of median absolute deviation (MAD) of ΔAUC in each dataset (MAD×3) and R^2^ = 0.8 cut-offs are indicated. Hits surpassing selection criteria, as well as PRDX6, are plotted in black. Thermal stability profiles for these hits are provided in Fig. S6. GluPho, glucose −6-phosphate dehydrogenase-6-phosphogluconolactonase (**H**) Quantification of lipid peroxidation damage using flow cytometric measurement of the fluorescent lipid peroxidation sensor BODIPY^581/591^ C11 following treatment of parasites with Darapladib (Dar) or Varespladib (Var) for 2h at trophozoite stage. (**I**) Ring to schizont progression and full-cycle growth curves following treatment with Darapladib (Dar) and α-tocopherol (Toco) co-treatment. Parasites were treated at ring stage. Data in (A-C, F and H-I) are means ± SD of three independent experiments, unpaired t-test.

To confirm that Darapladib specifically targets host PRDX6 to block blood stage parasite development, we exploited *P. yoelii* infected RBCs from transgenic *prdx6^-/-^* mice. We employed the *P. yoelii* YM lethal strain which readily invades mature RBCs similar to *P. falciparum* (Otsuki et al., 2009). The sequence of PRDX6 is highly conserved amongst mammalian species with 93% similarity between mouse and human PRDX6 (Leyens et al., 2003). Transgenic mice that are null for PRDX6 are viable, possibly because they compensate for loss of PRDX6 by increased expression of other antioxidant enzymes and higher glutathione levels (Feinstein, 2019; Melhem et al., 2017). Indeed, RBCs from *prdx6^-/-^* mice had significantly higher levels of peroxiredoxin 2 (PRDX2) compared to RBCs from WT mice (Fig. S5). It was, therefore, not surprising to find that growth of *P. yoelii* YM was similar in WT and *prdx6^-/-^* mice (Fig. 2E). This allowed us to assess the effect of Darapladib on *ex vivo* progression of *P. yoelii* YM from ring to schizont stages in RBCs from WT and *prdx6^-/-^* mice (Fig. 2F). Darapladib blocked *ex vivo* progression of the *P. yoelii* YM strain in WT mouse RBCs but had no effect on progression in RBCs from *prdx6^-/-^* mice up to a concentration of 1μM confirming that it specifically targets PRDX6 (Fig. 2F).

To identify possible off-targets of Darapladib in *P. falciparum* infected RBCs, we carried out a mass spectrometry based isothermal dose response cellular thermal shift assay (ITDR-CETSA) (Fig. 2G, Fig. S6) (Dziekan et al., 2019, 2020). Parasite lysates were exposed to varying concentrations of Darapladib, heat challenged and analysed for occurrence of drug-dose dependent changes in protein stability. Among 1782 parasite proteins and 277 human proteins detected in both biological replicates, no protein exhibited reproducible change in stability of ≥ 30% in the presence of ≤ 10μM Darapladib.

Some dose dependent changes in protein stability were observed in 10 proteins (Fig. S6) including in one essential *P. falciparum* enzyme, glucose-6-phosphate dehydrogenase-6-phosphogluconolactonase (*Pf*GluPho) (Fig. S6). However, these changes were either observed at high drug doses (>10μM) or were not reproducible in a second biological replicate and were therefore discounted. We did investigate further *Pf*GluPho, the only essential *P. falciparum* enzyme identified by IDTR-CETSA in one out of two biological replicates. Using both recombinant *Pf*GluPho as well as parasite lysate, we found that Darapladib did not inhibit the essential glucose-6-phosphate dehydrogenase (G6PD) activity of *Pf*GluPho in orthogonal enzymatic assays (Fig. S7). *Pf*GluPho was thus ruled out as an off-target of Darapladib. Human PRDX6 was identified in the ITDR-CETSA, but changes in thermal stability observed following Darapladib treatment were not statistically significant under assay conditions. Thus, we did not find any evidence for the presence of any relevant off-targets by ITDR-CETSA analysis, supporting PRDX6 as the exclusive target of Darapladib.

PRDX6 is a lipid peroxidation repair enzyme involved in protection of cells from oxidative damage. The fluorescent lipid peroxidation sensor BODIPY^581/591^ C11 was used to assess whether treatment of *P. falciparum*-infected RBCs with Darapladib resulted in increased lipid peroxidation damage (Fig. 2H). As expected, Darapladib treatment increased lipid peroxidation damage in *P. falciparum* blood stages in a dose-dependent manner (Fig. 2H). Increased lipid peroxidation damage was also confirmed independently by measurement of malondialdehyde (MDA) (Fig. S8). To further confirm the involvement of PRDX6 in lipid peroxidation repair, we co-treated parasites with Darapladib and the lipophilic antioxidant α-tocopherol (vitamin E), which prevents lipid peroxidation by scavenging radicals (Niki, 2014). Co-treatment with α-tocopherol rendered Darapladib ineffective and restored parasite progression and growth in a dose-dependent manner (Fig. 2I). These observations further demonstrate that host PRDX6 is essential for repair of lipid peroxidation damage during *P. falciparum* blood stage development.

### Inhibition of PRDX6 blocks vesicular transport of hemoglobin to FV

The localization of PRDX6 within HCV and inhibition of progression at trophozoite stage suggested that PRDX6 may play a role in HCCU and hemoglobin catabolism. We, therefore, tested the impact of Darapladib treatment on hemoglobin transport to the FV. For this, we co-treated parasites with Darapladib and the cysteine protease inhibitor E64 that blocks proteolytic digestion of hemoglobin leading to bloating of the FV (Lazarus et al., 2008). As expected, E64 treatment led to an enlarged FV due to accumulation of undigested hemoglobin, as observed by light microscopy (Fig. 3A and B) and transmission electron microscopy (Fig. 3C). In contrast, treatment with Darapladib, either alone or with E64, prevented bloating of the FV and resulted in parasites with a shrunken FV that contained little to no hemozoin suggesting that Darapladib acts upstream of hemoglobin digestion in the FV. (Fig. 3A and C). Moreover, multiple HCV were observed in the parasite cytosol upon treatment with Darapladib (Fig. 3C). Taken together, these observations suggest that PRDX6 is not involved in HCCU and HCV formation, but is essential for active transport of HCV to the FV.

**Figure 3.**
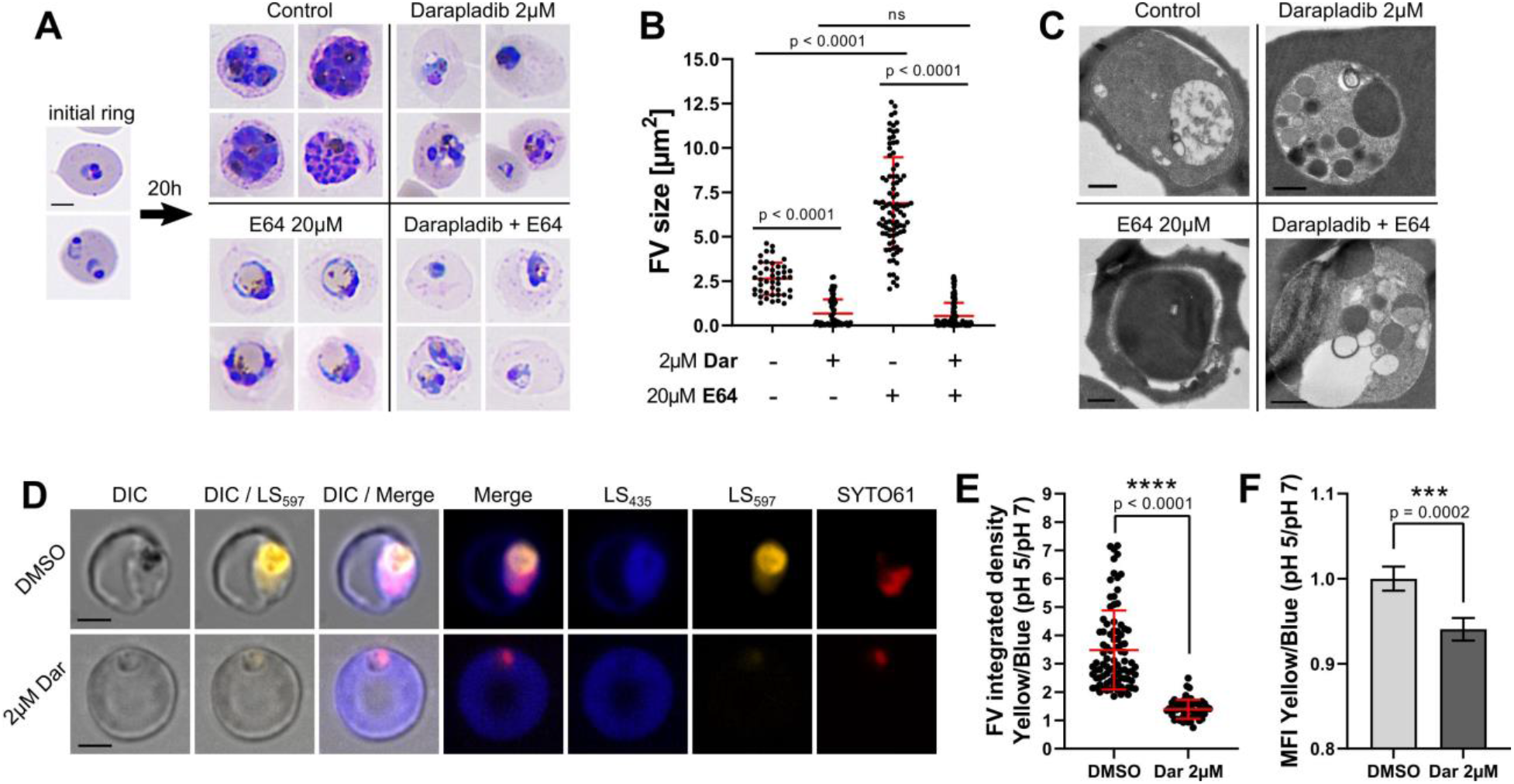
Inhibition of PRDX6 blocks vesicular transport of hemoglobin-containing vesicles (HCV) to the FV. (**A**) Light microscopy of *P. falciparum* blood stage cultures treated at the ring stage with Darapladib, E64 or Darapladib + E64 and incubated for 20h. Treatment of *P. falciparum* rings with E64 resulted in bloating of the FV with undigested hemoglobin. In contrast, treatment with either Darapladib alone or with E64 prevented E64-mediated bloating of the FV indicating a role of PRDX6 upstream of hemoglobin digestion. Scale bars: 5 μm. (**B**) Quantification of FV size observed in (A). (**C**) Transmission electron microscopy of *P. falciparum* blood stage cultures from (A). Darapladib arrested transport of HCV within the parasite cytosol. Scale bars: 500nm. (**D**) Fluorescence microscopy. Use of pH sensitive fluorescent probe to observe HCCU in *P. falciparum* blood stages following treatment with Darapladib (Dar). RBCs were preloaded with pH-sensitive LysoSensor Blue/Yellow, infected with *P. falciparum,* treated with Darapladib at the ring stage and imaged at the trophozoite stage. Darapladib treatment prevented transport of RBC cytosol (neutral pH, blue colour) to the FV (acidic, yellow colour). Scale bars: 5 μm. Dar, Darapladib; DIC, Differential Interference Contrast; LS_435_, LysoSensor at neutral pH; LS_597_, LysoSensor at acidic pH; SYTO61, DNA (nuclei). (**E**) Quantification of Yellow/Blue signal ratio within the FV observed by fluorescence microscopy in (D). (**F**) Flow cytometric measurement of Yellow/Blue signal in *P. falciparum*-infected RBCs treated with Darapladib as described in (D). Data are means ± SD of three independent experiments, unpaired t-tests.

To confirm that PRDX6 inhibition leads to disruption of delivery of host cell cytosol to the FV, we preloaded RBCs with the pH-dependent LysoSensor Blue/Yellow that emits blue fluorescence at neutral pH as found in the cytosol, and yellow fluorescence at acidic pH, as found in the FV. RBCs pre-loaded with LysoSensor Blue/Yellow were infected with *P. falciparum*, treated at ring stage and allowed to develop into trophozoites. Control parasites successfully internalized host cell cytosol and exhibited a bright yellow signal in the FV (Fig. 3D-F). Inhibition of PRDX6 with Darapladib reduced yellow fluorescence signal intensity and parasites appeared shrunken and pyknotic without visible hemozoin. Furthermore, treatment of *P. falciparum* rings with Darapladib significantly decreased the size of hemozoin crystals (Fig. S9). Taken together, these observations show that PRDX6 is essential for the transport of hemoglobin to the FV.

### Co-treatment of artemisinin-resistant parasites with artemisinin and Darapladib synergistically reduces parasite survival

Resistance against the frontline antimalarial drug artemisinin (ART) is a major threat for malaria control and elimination efforts. Parasites attain ART-resistance through reduced hemoglobin uptake and digestion, which lowers oxidative stress levels and precludes activation of ART (Birnbaum et al., 2020; Xie et al., 2020). As inhibition of PRDX6 by Darapladib raises oxidative stress levels, we investigated whether co-treatment with Darapladib and ART could restore ART-susceptibility in ring stage survival assays (RSA) (Witkowski et al., 2013). RSA of *P. falciparum* ring stages co-treated with Darapladib and Dihydroartemisinin (DHA), one of the ART derivatives used as first-line antimalarials, synergistically reduced survival of NF54 Kelch13^C580Y^ with complete reversal of ART-resistance with 1 μM Darapladib (Fig. 4A and Table S2). The ART-resistant field isolate from Cambodia, *P. falciparum* 3815 (Ariey et al., 2014), was also susceptible to co-treatment with Darapladib and DHA in the RSA (Fig. 4A and Table S2). Both these parasites have the Kelch13^C580Y^ mutation, but in different genetic backgrounds. Survival of the highly resistant field isolate 3601 carrying the Kelch13^R539T^ mutation also decreased following Darapladib co-treatment with ART, although the reduction was less compared to the NF54 Kelch13^C580Y^ and 3815 strains (Fig. 4A and Table S2). However, Darapladib effectively blocked blood stage growth of all tested strains with both Kelch13^R539T^ and Kelch13^C580Y^ mutations following continuous treatment during the complete blood stage cycle (Fig. 4B). Summed up, Darapladib could thus both reduce survival of ART-resistant parasites when combined with artemisinin as well as independently block parasite growth of ART-resistant *P. falciparum* strains.

**Figure 4.**
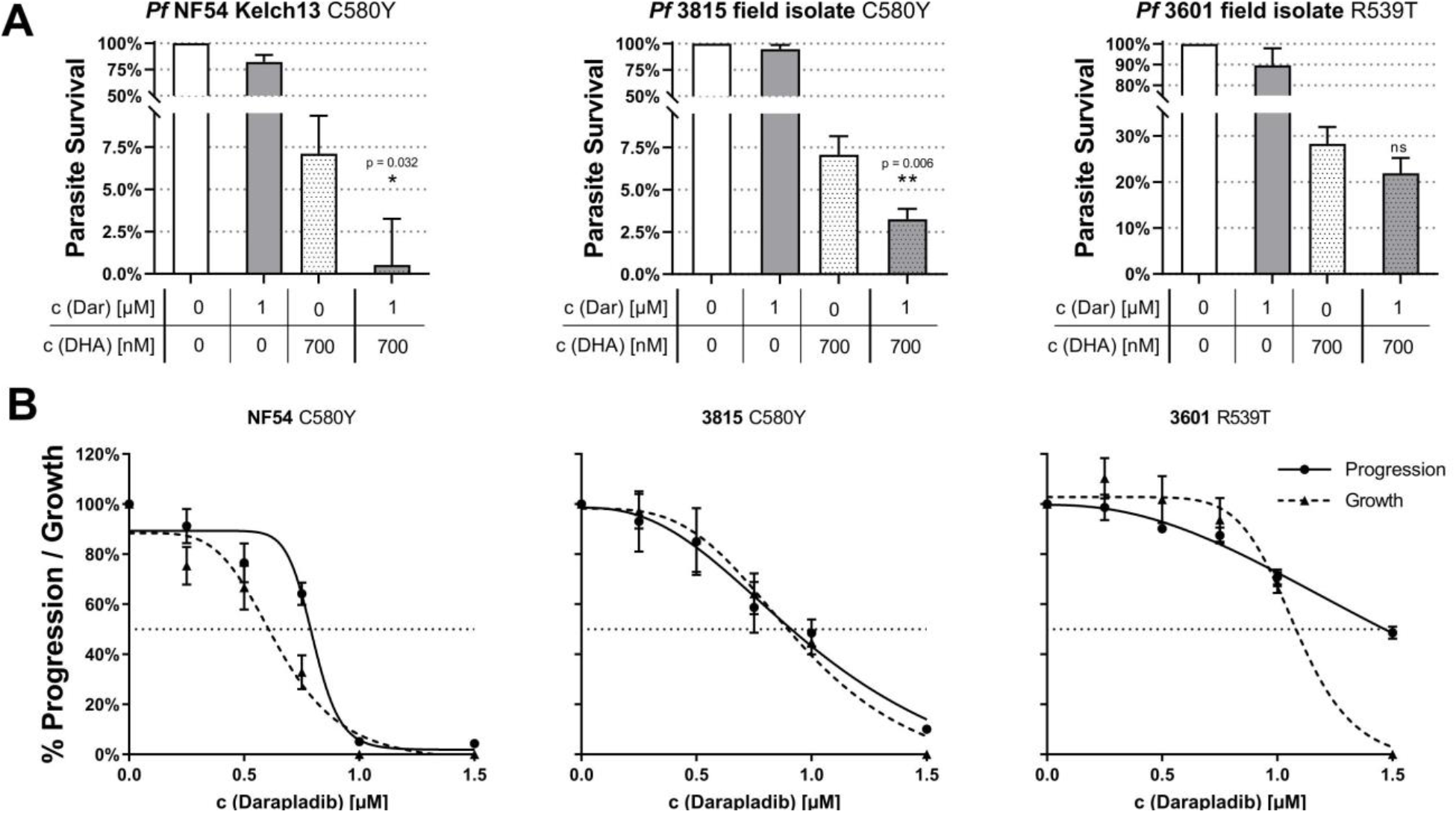
Co-treatment of ART-resistant parasites with artemisinin and Darapladib synergistically reduces parasite survival. (**A**) Ring stage survival assays with Darapladib (Dar) and Dihydroartemisinin (DHA) against ART-resistant *P. falciparum* strains. For detailed data regarding synergism cf. Table S2. (**B**) Growth/Progression assay with continuous Darapladib treatment against ART-resistant *P. falciparum* strains. Data are means ± SD of three independent experiments, unpaired t-tests.

## DISCUSSION

The import and proteolytic degradation of hemoglobin during *P. falciparum* blood stage growth creates significant oxidative stress that can cause oxidative damage to both host and parasite lipid membranes. Interestingly, *P. falciparum* has been shown to internalize host antioxidant enzymes such as catalase, superoxide dismutase and PRDX2 by unknown mechanisms during blood stage infection (Foth et al., 2011; Koncarevic et al., 2009). However, these enzymes exclusively detoxify small free reactive oxygen species (ROS) and are not involved in lipid peroxidation repair. Mechanisms for the management and repair of lipid peroxidation damage during *P. falciparum* blood stage growth are unclear and no lipid peroxidation repair enzymes have been identified in the parasite proteome (Flammersfeld et al., 2018; Fu et al., 2009). Here, we demonstrate that *P. falciparum* imports the host lipid peroxidation repair enzyme PRDX6 during the blood stage allowing it to repair lipid peroxidation damage. Moreover, we show that PRDX6 also plays an important role in transport of hemoglobin-containing vesicles (HCV) to the food vacuole (FV).

Host cell cytosol uptake (HCCU) is a key mechanism for the import of hemoglobin during blood stage growth of malaria parasites. Research on HCCU has so far focused on uptake of hemoglobin, but has neglected the potential import and role of other cytosolic RBC proteins. Our study shows for the first time that a host lipid peroxidation repair enzyme, PRDX6, is internalized along with hemoglobin during HCCU. Early cytostomes and HCVs predominantly contain host cytosolic proteins. Thus, it is not surprising that antioxidant protection in these vesicles might primarily be facilitated by host cytosolic antioxidant enzymes. The observation that cytostomes and HCVs have higher concentrations of hemoglobin and PRDX6 than the RBC cytosol implies selective vesicular loading mechanisms, which remain to be identified.

Based on our observations, we propose a model where PRDX6 is essential for lipid peroxidation repair in nascent HCVs as shown in the Graphical Abstract. Inhibition of PRDX6 leads to increase in lipid peroxidation damage in HCVs due to lack of lipid repair, which appears to disrupt active transport of HCVs towards the FV. As a result, HCVs accumulate in trophozoite stages, as observed by electron microscopy of Darapladib treated parasites. The mechanistic understanding of active transport of HCVs in *Plasmodium spp*. is unfortunately limited. In mammalian cells, vesicular transport is facilitated by actin-myosin motor proteins (Lee Sweeney and Holzbaur, 2018). Myosin binds to myosin receptors on the vesicle surface to anchor the motor and enable movement. Elevated lipid peroxidation damage may disturb the localization or functioning of such receptors in the HCV membrane disrupting normal attachment of HCVs to their respective motor proteins leading to disruption of active transport. This notion is supported by several studies that implicate actin in HCV transport (Lazarus et al., 2008; Smythe et al., 2008). Conditional inactivation of the parasite protein *Pf*VPS45 causes a similar phenotype of arrested HCVs in the parasite cytosol suggesting a functional role for *Pf*VPS45 in vesicular transport to the FV (Jonscher et al., 2019).

Darapladib was originally developed as a selective inhibitor of extracellular lipoprotein-associated PLA_2_ (Lp-PLA_2_) (Blackie et al., 2003). However, Lp-PLA_2_ is absent in RBCs and also not internalized by *P. falciparum* infected RBCs from blood serum (Gautier et al., 2018; Tougan et al., 2020; Zhou et al., 2011). Darapladib was shown to bind PRDX6 using ABPP and blocks the PLA_2_ activity of PRDX6. The on-target specificity of Darapladib was demonstrated using RBCs from *prdx6^-/-^* mice that lack PRDX6. In the absence of PRDX6 in RBCs from *prdx6^-/-^* mice, Darapladib loses its inhibitory effect on progression of *P. yoelii* from rings to schizonts. The observation that Darapladib inhibits blood stage progression of both *P. falciparum* and *P. yoelii* suggests a common mechanism for lipid peroxidation damage repair that is essential for parasite survival across different *Plasmodium* species. The specificity of Darapladib for PRDX6 was furthermore validated using ITDR-CETSA, which demonstrated that no other host or parasite protein is targeted by Darapladib. However, due to the relatively high IC_50_ of Darapladib for inhibition of blood stage growth (≈ 0.5μM), Darapladib itself is not suitable for use as an antimalarial drug candidate. Also, due to the low bioavailability of Darapladib, inhibition of PRDX6 by Darapladib cannot be tested in a clinical *in vivo* malaria model (Dave et al., 2014). Since no high affinity inhibitor of PRDX6 is known, improved inhibitors with a higher affinity to PRDX6 need to be identified and could serve as promising antimalarial drug candidates.

Resistance to artemisinin (ART), one of the major threats to malaria control efforts, which arose in *P. falciparum* strains in Southeast Asia has now been identified on the African continent (Balikagala et al., 2021; Uwimana et al., 2021). Strategies to combat artemisinin resistance are urgently needed. We found that co-treatment with ART and Darapladib synergistically reduced survival of ART-resistant parasites. PRDX6 inhibitors could thus play a pivotal role in restoring the efficacy of artemisinin against ART-resistant strains. In addition, a PRDX6 inhibitor will independently block growth of *P. falciparum* blood stages by inhibiting transport of hemoglobin to the FV. An ART-based combination therapy involving artemisinin together with a PRDX6-targeting partner drug would be effective in two ways: Firstly, it could help to synergistically reduce ART-resistant parasite survival during early ring stage, restoring ART-susceptibility. Secondly, it would independently inhibit blood stage growth of *P. falciparum* isolates including ART-resistant strains.

The identification of PRDX6, a host RBC enzyme that is essential for *P. falciparum* blood stage growth, allows the development of completely novel strategies for pharmacological treatment and control of malaria. Approaches that target host enzymes for development of anti-malarial drugs are attractive because parasites cannot mutate the target gene to attain resistance (Wei et al., 2021). Thus, inhibitors that target host enzymes such as PRDX6 are less likely to face the problem of drug resistance and could play a critical role in malaria eradication efforts in the long term.

## AUTHOR CONTRIBUTIONS

Conceptualization: MPW, CEC

Methodology: MPW, LT, CEC

Investigation: MPW, PF, OG, JMD, IB

Funding acquisition: CEC, MPW

Project administration: CEC

Supervision: CEC

Resources: DM, JCB, RA, CH, SR, ZB, ABF, SIF

Writing: MPW, CEC

## ACKNOWLEDGMENTS

We thank Gordon Langsley for fruitful discussions and critical inputs. We also thank the electron microscopy Ultrastructural BioImaging (UBI) platform, Institut Pasteur, Paris, for providing ready access to the electron microscopy facility. The authors are grateful to the Central Animal Facility of Institut Pasteur for providing support with animal importation, care, housing and breeding. We thank the Image Analysis Hub (IAH), Institut Pasteur, Paris, for their help with image analysis. MPW was part of the Pasteur - Paris University (PPU) International PhD Program and European Union’s Horizon 2020 research and innovation program under the Marie Sklodowska-Curie grant agreement No 665807. This work was funded by the following grants:

Fondation pour la Recherche Médicale grant (FDT202001010791) to MPW

Agence Nationale de la Recherche grant (ANR-17-CE15-0010) to CEC

Pasteur Innov grant from Institut Pasteur to CEC

Laboratoire d’Excellence (LabEx) “French Parasitology Alliance For Health Care” (ANR-11-LABX-0024-PARAFRAP) to CEC and DM.

Laboratoire d’Excellence ‘Integrative Biology of Emerging Infectious Diseases’ (grant no. ANR-10-LABX-62-IBEID) to PF

NTU Presidential Postdoctoral Fellowship Grant (NTU/PPF/2019) to JMD

LOEWE Center DRUID (Projects B3) within the Hessian Excellence Program to IB

## DECLARATION OF INTERESTS

ABF and SIF are shareholders in Peroxitech LLC, USA a company that is developing a peptide based PRDX6 inhibitor as a therapeutic agent.

## RESOURCE AVAILABILITY

### Lead Contact For Reagent And Resource Sharing

Further information and requests for reagents should be directed to and will be fulfilled by the Lead Contact, Chetan Chitnis.

### Materials availability

This study did not generate new unique reagents.

### Data and code availability

The mass spectrometry proteomics data have been deposited to the ProteomeXchange Consortium via the JPOSTrepo partner repository with the dataset identifier PXD029803. Microscopy data reported in this paper will be shared by the lead contact upon request. This study did not generate new code. Any additional information required to reanalyze the data reported in this paper is available from the lead contact upon request.

## EXPERIMENTAL MODEL AND SUBJECT DETAILS

### *Plasmodium falciparum* parasites

All *Plasmodium falciparum* strains were cultured at a final hematocrit of 2–4% under mixed gas atmosphere (5% O_2_, 5% CO_2_ and 90% N_2_) at 37°C. *P. falciparum* 3D7 (Walliker et al., 1987) was cultured in RPMI-1640 medium with 0.5% Albumax (complete RPMI, cRPMI) using O^+^ human erythrocytes (ICAReB, Institut Pasteur Paris, France). ART-resistant parasite strains *P. falciparum* NF54 C580Y (Ariey et al., 2014) and the field isolates Cambodia 3815 and 3601 (Ariey et al., 2014) were cultured in RPMI supplemented with 0.5% Albumax and 2.5% AB^+^ human serum (EFS, Rungis, France) using O^+^ human erythrocytes.

### Red blood cells

O^+^ human erythrocytes in citrate buffer were obtained commercially from ICAReB, Paris, France.

### *In vivo* infection of mice with *P. yoelii*

*prdx6^-/-^* mice were generated from C57/BL6JRj mice in the lab of Aron B. Fisher, University of Pennsylvania, USA as described earlier and are available under the accession number B6.129-*Prdx6^tm1Abf^*/Mmjax (Feinstein, 2019). Animals were housed in the Institut Pasteur animal facilities accredited by the French Ministry of Agriculture for performing experiments on live rodents. Work on animals was performed in compliance with French and European regulations on care and protection of laboratory animals (EC Directive 2010/63, French Law 2013-118, February 6th, 2013). All experiments were approved by the Ethics Committee #89 and registered under the reference 01324.02 and dap180040. At the time of the experiment, both *prdx6^-/-^* mice and WT mice were about 44 weeks of age and littermates of both sexes were randomly assigned to experimental groups.

*prdx6^-/-^* mice and age-matched control C57/BL6JRj WT mice (Janvier Labs, France) were infected with the *P. yoelii* YM lethal strain (Mwakingwe et al., 2009) via the tail vein for an initial parasitemia of 0.002%. Parasitemia was measured by Giemsa-stained blood smears and by flow cytometry. For this, *P. yoelii* YM cells were labelled with 5× SYBR Green I and 1:100 α-mouse CD71-APC (eBioscience, UK) for 30min at 37°C. To specifically assess intraerythrocytic growth, CD71^+^ reticulocytes were excluded. Infected mature RBCs were identified as SYBR Green^+^ and CD71^-^ cells.

## METHOD DETAILS

### Lysis with Saponin and sample preparation

Late trophozoites and early schizonts from synchronous *P. falciparum* 3D7 cultures were purified with a Percoll gradient. The resulting parasite pellet was resuspended in PBS. RBC membranes were lysed by exposure to 0.01% Saponin (Sigma, Stock 1% in H_2_O) for 5 min at RT. Complete hemolysis was confirmed by Giemsa-stained blood smears. After washing two times with PBS, the hemolysed pellet was either assessed by mass spectrometry (cf. cellular thermal shift assay) for immunoblotting, it was resuspended in 1× Laemmli SDS sample buffer (BioRad) with 200mM DTT (Sigma) and boiled at 95°C for 10 min.

### SDS-PAGE and Immunoblotting

Samples were separated on SDS-polyacrylamide gel and proteins were transferred onto a nitrocellulose membrane (GE healthcare) using a wet transfer system (BioRad). The membrane was blocked in PBS with 5% skimmed milk overnight at 4°C with continuous agitation. Following this, the membranes were incubated with mouse monoclonal antibody against human PRDX6 (α-hPRDX6 1A11 (1:2000), Santa Cruz), rabbit antisera raised against *P. falciparum* nucleosome assembly protein L (α-*Pf*NAPL 1:500) (Singh et al., 2010) and goat antisera against human hemoglobin (α-hHb HRP 1:5000, Bethyl) in PBS with 2.5% skimmed milk and 0.05% Tween 20 for 2h at RT. After washing, blots with α-PRDX6 and α-*Pf*NAPL were incubated with respective α-mouse and α-rabbit horse radish peroxidase (HRP)-coupled secondary antibodies (1:5000, Promega) for 1h at RT. Membranes were developed with enhanced chemiluminescence (ECL) substrate (ThermoFisher Scientific) and imaged on an Amersham Imager 600 (GE Healthcare). Band intensity was calculated using ImageJ software (NIH, Bethesda, USA) (Schneider et al., 2012).

### Immuno-fluorescence microscopy

*P. falciparum* 3D7 was cultured as described above. 1mL of a 4% hematocrit, high parasitemia culture was pelleted by centrifugation (1.5 g, 3min, RT) and resuspended in freshly prepared PBS containing 4% paraformaldehyde (Electron Microscopy Sciences) and 0.0075% gluteraldehyde (Sigma) and fixed for 30 min at RT. After washing in PBS once, fixed parasites were permeabilized with 0.1% Triton X-100 in PBS for 10 min at RT. Cells were washed in PBS again and blocked for 30 min in PBS with 2.5% BSA. Antibody dilutions were prepared in PBS with 2.5% BSA. After blocking, parasites were incubated with mouse α-hPRDX6 1A11 (1:500, Santa Cruz) for 1h at RT. Parasites were washed twice in PBS and incubated with α-mouse AlexaFluor 488-coupled secondary antibody (1:2000, Invitrogen) and Hoechst 33342 (2 μM final) (Invitrogen), washed again twice in PBS and resuspended in 30 μL PBS. From this suspension, 4 μL was transferred onto a glass slide and covered with a glass coverslip. Samples were examined under a Deltavision Elite high resolution fluorescence microscope (GE Healthcare Lifesciences) using DAPI, FITC and differential interference contrast (DIC) channels. Hemozoin crystal size was measured in DIC images using ImageJ software (NIH, Bethesda, USA).

### Transmission electron microscopy

#### Fixation

1 mL of *P. falciparum* 3D7 culture was pelleted by centrifugation (1500g, 3 min, RT) and resuspended in freshly prepared PBS containing 4% EM-grade para-formaldehyde (PFA, Electron Microscopy Sciences) and 0.1% EM-grade GA (Sigma) and fixed for 1 h on ice. After washing in PBS once, fixed parasites were stored in PBS with 1% PFA at 4°C.

#### Immuno-TEM

Fixed parasites were washed three times in PBS followed by incubation for 15min in 50mM NH4Cl at RT. Parasites were resuspended in PBS with 10% gelatin (Sigma) and incubated for 10 min at 37°C. The samples were allowed to solidify on ice and were infiltrated with 2.3M sucrose at 4°C overnight, mounted onto sample pins and frozen in liquid nitrogen. Subsequently, the samples were cryo-sectioned (60 nM thickness) using a FC6/UC6 cryo-ultramicrotome (Leica) and a 35° diamond knife (Diatome). The sections were picked up using a 1:1 mixture of 2% methyl cellulose (Sigma) and 2.3M sucrose. Samples were quenched with 50 mM NH_4_Cl, blocked in PBS with 1% BSA and immunolabeled with α-hPRXD6 1A11 (SantaCruz, 1:20) IgG1a monoclonal antibody in PBS with 1% BSA followed by a bridge step labelling with a polyclonal Rabbit Anti-Mouse Immunoglobulins (Dako, 1:50) and protein A-gold 10 nm (UMC Utrecht, 1:50) treatment for 15 min. Finally, samples were fixed with 1% glutaraldehyde in PBS for 5 min. The sections were thawed and stained/embedded in 4% uranylacetate/2% methyl cellulose mixture (1:9). Images were recorded with a Tecnai Spirit transmission electron microscope at 120 kV with bottom-mounted EAGLE 4Kx4K camera.

#### EPON resin embedding

Fixed parasites were treated again with 2.5% gluteraldehyde (Sigma) in PBS overnight, washed three times in PBS and mordanted with tannic acid 1% in PBS for 30min at RT. Cells were then washed three times in PBS, postfixed with 1% OsO_4_ for 1.5 h and washed three times with distilled water and stored overnight at 4°C. Samples were dehydrated 15 min in a graded series of ethanol (25%, 50%, 75%, 95%, 3× 100%). Samples were incubated 1×15min, then 2×10min in propylene oxide and embedded in EMbed-812 epoxy resin (EMS; EPON/Propylene Oxide: r = 25/75 for 2 h; r = 50/50 overnight; r = 75/25 all the day and overnight with the tubes open under the chemical hood). Samples were then embedded 3× 2h at RT in pure EPON and subjected to heat polymerization for 5 days at 60°C. Thin sections were cut with a Leica Ultramicrotome Ultracut UC7’ sections (60 nm), stained with uranyl acetate and lead citrate. Images were recorded with a Tecnai Spirit transmission electron microscope at 120 kV with bottom-mounted EAGLE 4Kx4K camera (FEI-Thermofisher).

### Recombinant expression of human PRDX6

A synthetic gene encoding human PRDX6 was tagged with a 6x-His tag, codon-optimized for expression in *E. coli* was synthesized and cloned into the *E. coli* expression vector pET28a (GenScript, Piscataway, USA). The plasmid vector was transformed into *E. coli* BL21 (DE3) competent cells (Sigma). A single colony of transformed bacteria was picked from a LB-agar plate with Kanamycin (50 μg/mL) and grown in LB medium with Kanamycin at 37°C to an optical density of OD_600_ of 0.6. Expression of recombinant PRDX6 was induced with 1mM isopropyl β-d-thiogalactopyranoside (IPTG, Sigma). The culture was incubated for 20h at 18°C, bacteria were harvested by centrifugation (6000 g, 10 min, 4°C) and resuspended in lysis buffer (50 mM Tris/HCl at pH 8, 150 mM NaCl, 2 mM β-mercaptoethanol). After sonication and removal of cell debris by centrifugation, the supernatant was loaded onto a HisTrap FF affinity column (GE Healthcare) and equilibrated with lysis buffer (see above). Recombinant PRDX6 was eluted with lysis buffer supplemented with 250 mM imidazole (Sigma). Subsequently, recombinant PRDX6 was purified by gel filtration chromatography on a Superdex 200 16/600 column (Sigma) equilibrated with 20mM HEPES at pH 7.0, 2mM EDTA and 1mM DTT. The yield from a 1L initial bacterial culture was about 12.5 mg of purified recombinant PRDX6.

### Activity-based protein profiling of PRDX6

Recombinant human PRDX6 was diluted to 10 mg/mL in PBS. Aliquots of 50 μL were incubated for 5 min at RT with different concentrations of Darapladib (SelleckChem, USA), Varespladib (Cayman Chemical, USA), MAFP (Cayman Chemical, USA), ATK (Cayman Chemical, USA), P11 (Cayman Chemical, USA) or DMSO. TAMRA-fluorophosphonate (TAMRA-FP, Invitrogen, Thermo Fisher, USA) was solved in DMSO (100 μM), added to the sample (2 μM final) and incubated for 30min at 37°C. After 30min, the reaction was quenched by addition of 15 μL 4×Laemmli sample buffer and 7.5 μL 1M DTT. The sample was heated to 95°C for 10 min and separated by SDS-PAGE on a 12% SDS polyacrylamide gel. After electrophoresis, the gel was imaged on a ChemiDoc MP imager (BioRad, USA) in the Cy3 channel.

### PRDX6 phosphorylation and PLA_2_ activity assay

PRDX6 was phosphorylated according to the method described by Wu et al. (Wu et al., 2009). Recombinant human PRDX6 (150 μg/mL final) and active MAPK (ERK2, R&D Systems, 10 μg/mL final) were added to 30 μL of a phosphorylation buffer containing 50 mM Tris/HCl pH 7.5, 20 μM EGTA, 10 mM MgCl_2_ and 2 mM Mg-ATP (Sigma) and were incubated for 90 min at 30°C.

Measurement of PLA_2_ activity of phosphorylated PRDX6 (pPRDX6) was based on the enzymatic PLA_2_ assay described by Wu et al. (Wu et al., 2009) and the rapid free fatty acid extraction method by Katsumata et al. (Katsumata et al., 1986). Liposomes consisting of DPPC/egg yolk PC/egg yolk PG/cholesterol (6:3:1.2:0.95, DPPC from Avanti Polar Lipids, all other Sigma) with 0.6 μCi tracer [2-palmitoyl-1-^14^C]-dipalmitoyl phosphatidylcholine (^14^C-DPPC, American Radiolabeled Chemicals) were prepared by freezing/thawing three times in liquid nitrogen. 100 μL diluted phosphorylated PRDX6 (2.5 μg/mL in PLA_2_ assay buffer 40 mM NaOAc, 5 mM EDTA, pH 5), or blank PLA_2_ assay buffer was added to 800 μL PLA_2_ assay buffer containing different concentrations of Darapladib in DMSO. After addition of 100 μL liposome preparation, the samples were incubated for 2 h at 37°C. The enzymatic reaction was terminated by addition of 200 μL 5% Triton X-100 in PLA_2_ assay buffer. The product of the PLA_2_ enzymatic reaction, free ^14^C-palmitic acid, was extracted by addition of 10 mL hexane with 0.1% acetic acid (v/v) and 200 mg of anhydrous Na_2_SO_4_ (Tong et al., 1998) and subsequent vortexing for 20s. 3 mL of the hexane layer was added to 10 mL UltimaGold liquid scintillation cocktail (PerkinElmer, Waltham, USA) and measured on a TriCarb 2800TR liquid scintillation counter (PerkinElmer, Waltham, USA).

### Parasite progression and growth assay

#### General assay description

To assess the effect of inhibitors on the ring to schizont progression, and growth from ring stage to next generation ring stage, a tightly Percoll-synchronised culture with about 2% parasitemia and 2% hematocrit was treated with different inhibitors at ring stage (14-20 hpi). The inhibitors were present throughout the assay. The concentration of the solvent was kept equal in all samples in a given experiment and maintained below 0.1% to avoid toxic effects. For each condition, a triplicate set of 1 mL cultures was added into wells of a 24-well plate and kept at 37°C and 5% O_2_, 5% CO_2_ throughout the assay. The initial ring stage culture was scored by flow cytometry as described below and by examination of Giemsa-stained blood smears. After 20h, the samples were measured again to assess progression to schizont stage. Further 24h later, the overall growth and development of next generation rings was measured. Flow cytometry data was analyzed with the FlowJo 10.8 software (FlowJo LLC). After exclusion of doublets, a gate for uninfected and infected RBCs based on the size of the cells in the FSC-A channel was set up to exclude debris and merozoites. Finally, gates for rings, trophozoites and multiply infected RBCs, and schizonts were set up. The total parasitemia was defined as the sum of the aforementioned gates. Each experiment was performed with three biological replicates. Relative rate of progression was calculated as the fraction of schizonts in treated and control samples (T/C). Relative growth was calculated as the ratio of parasitemia in the treated and control samples after subtraction of the initial parasitemia (T-I/C-I).

#### Labelling of P. falciparum for flow cytometry

Cultures were diluted to 0.2% hematocrit and transferred to a 96-well round bottom plate. 5 μL of 200× SYBR Green I (Lonza, Switzerland) was added to 200 μL of the diluted culture for a final concentration of 5× SYBR Green I. The plate was then incubated for 30 min at 37°C in the dark. Samples were measured on a calibrated MACSquant flow cytometer on the forward scatter (FSC), side scatter (SSC) and B1 fluorescent channel (B1: λ_Ex_ 488 nm and λ_Em_ 525/50 nm). Data was processed with the FlowJo 10.8 Software (FlowJo, LLC).

### Infection of mice with *P. yoelii*

*prdx6^-/-^* mice and age-matched control C57/BL6JRj WT mice (Janvier Labs, France) were infected with the *P. yoelii* YM lethal strain (Mwakingwe et al., 2009) via the tail vein for an initial parasitemia of 0.002%. Parasitemia was measured by Giemsa-stained blood smears and by flow cytometry. *P. yoelii* YM cells were labelled with 5× SYBR Green I and 1:100 α-mouse CD71-APC (eBioscience, UK) for 30min at 37°C. To specifically assess intraerythrocytic growth, CD71 ^+^ reticulocytes were excluded. Infected mature RBCs were identified as SYBR Green^+^ and CD71^-^ cells.

### *Ex vivo P. yoelii* YM progression assay

*prdx6^-/-^* mice and age-matched control C57/BL6JRj WT mice were infected with *P. yoelii* YM as described above. When the parasitemia in the mice reached about 20%, 20μL blood was drawn from the chin and collected in heparinized tubes. 15μL of the blood was diluted in 8 mL of RPMI-1640 medium with 0.5% Albumax (cRPMI) and added in 1mL aliquots into a 24-well plate. The initial amount of schizonts was measured by Giemsa-stained blood smears and flow cytometry by labelling the parasites with 5× SYBR Green I and 1:100 α-mouse CD71-APC for 30min at 37°C. Infected mature erythrocytes were identified as SYBR Green^+^ and CD71^-^ cells. CD71^+^ reticulocytes were excluded by gating. Cells were treated with Darapladib (0.5 and 1μM) or DMSO and incubated for 16 h at 37°C, 5% O_2_ and 5% CO_2_ to allow progression of parasites to schizont stage. After 16 h, schizont development in RBCs was assessed by Giemsa-stained blood smears and flow cytometry as described above.

### MS-IDTR-CETSA and analysis

The assay was carried out in two biological replicates, as described with minor modifications (Dziekan et al., 2020). In brief, lysate from saponin-liberated mature (32±6 hpi) *P. falciparum* parasites was incubated for 3min with different concentrations (1.5nM– 100μM) of Darapladib (MedChemExpress) or the DMSO control and subsequently heated for 3min to 50°C or 55°C to denature unstable protein subsets, or incubated at 37°C. Soluble protein fractions were isolated by centrifugation and analysed by quantitative mass spectrometry. Data analysis was carried out in R environment using mineCETSA package. Resulting protein stability profiles under thermal challenge conditions (i.e. > 37°C) were evaluated based on observed change in protein stability [reading delta Area Under the Curve (ΔAUC) relative to the non-denaturing (37°C) condition, the Fold Change (FC) in protein abundance relative to vehicle control treated sample], as well as protein stability profile adherence to expected sigmoidal shape of dose response (R^2^). The criteria used for identification of drug-dose dependent changes in protein stability are characterized by ΔAUC surpassing two or three times Median Absolute Deviation (MAD) for the dataset, FC>0.3 and R^2^>0.8. The mass spectrometry proteomics data have been deposited to the ProteomeXchange Consortium via the JPOSTrepo partner repository with the dataset identifier PXD029803.

### Recombinant expression of *P. falciparum* GluPho

Recombinant N-terminal His6-tagged *P. falciparum* glucose-6-phosphate dehydrogenase-6-phosphogluconolactonase (*Pf*GluPho) was produced according to Jortzik et al. (Jortzik et al., 2011) in the vector pQE30 in *E. coli* M15 cells (Qiagen) with pRAREII (Novagen).

### Glucose-6-phosphate dehydrogenase activity assay

Glucose-6-phosphate dehydrogenase (G6PD) activity of recombinant *Pf*GluPho was measured by monitoring the reduction of NADP^+^ to NADPH at ex340/em460 nm in a 96-well formate as described before (Preuss et al., 2012). Recombinant *Pf*GluPho (0.4 μg/mL) was added to a reaction mixture containing 0.05 M Tris (pH 7.5), 3.3 mM MgCl_2_, 0.005 % Tween20, 1 mg/mL BSA and the substrates NADP^+^ (20 μM, Cayman) and glucose 6-phosphate (G6P, 25 μM, Sigma) close to the K*_M_*. Linear reaction curves were monitored over 10 min, and IC_50_ values calculated with GraphPad Prism. Samples without glucose-6-phosphate (G6P) were used to control the specificity of the reaction.

Parasite lysate was prepared by sonication of Saponin (0.01%) liberated trophozoite stage *P. falciparum* 3D7 parasites. Briefly, *P. falciparum* lysate was added to a reaction mixture containing 0.1 M Tris/HCl (pH 8.0), 10mM MgCl_2_, 0.5mM EDTA and 200μM NADP^+^. 200μM of G6P were added as a substrate and the enzyme activity was measured on a Tecan infinite M1000 Pro plate reader at 340nm. To correct for background activity in the parasite lysate and to assess specificity of the reaction, samples without G6P substrate were measured.

### Lipid peroxidation measurement with BODIPY^581/591^ C11

To measure oxidative stress, parasites were labelled with the fluorescent lipid peroxidation sensor BODIPY^581/591^ C11 (Invitrogen, Thermo Fisher, USA) based on the method described by Fu et al. (Fu et al., 2009). A *P. falciparum* 3D7 culture was highly synchronized via repeated Percoll gradient purifications. Parasites were treated with 0.5 μM or 2 μM Darapladib, 10 μM Varespladib or DMSO for 2h at late trophozoite/early schizont stage (38–42 hpi). After treatment for 2 h, parasites were pelleted by centrifugation and resuspended in labelling solution (5 μM BODIPY^581/591^ C11, 10 μM Hoechst 33342 in cRPMI) and incubated for 1 h at 37°C, 5% O_2_, 5% CO_2_. Parasites were measured on a MACSquant flow cytometer: BODIPY^581/591^ C11 reduced, λ_Ex_ 488nm, λ_Em_ 614/50nm, “Red”; BODIPY^581/591^ C11 oxidised λ_Ex_ 488nm, λ_Em_ 525/50 nm, “Green”; Hoechst 33342, λ_Ex_ 405nm, λ_Em_ 450/50 nm. Data was analyzed using FlowJo 10 (FlowJo, LLC). After exclusion of doublets, debris and merozoites by size, infected RBCs were gated for by selecting Hoechst 33342 positive cells. The MFI of “Red” and “Green” channels of infected RBCs was calculated. Lipid peroxidation was calculated as MFI_Green_/(MFI_Green_ + MFI_Red_). Relative lipid peroxidation was calculated as the ratio of lipid peroxidation in treated and control parasites (T/C).

### TBARS assay

Detection of thiobarbituric acid reactive substances (TBARS), such as malondialdehyde (MDA), was adapted from the method published by Ohkawa et al. (Ohkawa et al., 1978). To quench auto-oxidation, butylated hydroxytoluene (BHT, Sigma) was added to the reaction (Pikul and Leszczynski, 1986). A 0.8% (w/v) aqueous solution of 2-thiobarbituric acid (TBA, Sigma) was prepared by heating the solution to 50°C for 2h. A 20% parasitemia, 4% hematocrit *P. falciparum* 3D7 culture, or a 4% hematocrit RBC suspension were treated with 0.5μM or 2μM Darapladib, or DMSO for 8h at 37°C, 5% O_2_, 5% CO_2_. Afterwards, 5mL of each culture was pelleted by centrifugation. The pellet was resuspended in a 15 mL reaction tube with 1.5 mL 20% acetic acid pH 3.5 (v/v, Sigma), 200μL 10% SDS (Sigma), 100μL of 0.1% (w/v) BHT in EtOH, 1.5mL of 0.8% TBA in H_2_O and 600 μL H_2_O for a total reaction volume of 4 mL. The samples were heated for 1 h in vigorously boiling water. To stop the reaction, the reaction mix was cooled down in an ice bath. The pink colored reaction product was extracted by addition of 5 mL 1 -butanol (Sigma), vortexing and centrifugation (2000 g, 10min, RT). 150μL of the butane layer was transferred into a black-walled 96-well plate and fluorescence (λ_Ex_ 530 nm, λ_Em_ 550 nm) was measured on a Tecan Infinite M1000 Pro plate reader.

### Host cell cytosol uptake assays

#### Host cell cytosol uptake assay with E64

A synchronous *P. falciparum* 3D7 culture at ring stage was treated with 2 μM Darapladib, 20 μM E64, a combination of both, or DMSO as control and incubated overnight at 37°C, 5% O_2_, 5% CO_2_ until control parasites progressed to schizonts. Cells were observed by light microscopy of Giemsa-stained blood smears and transmission electron microscopy of EPON resin embedded parasites as described above.

#### Host cell cytosol uptake assay with LysoSensor Y/B

RBCs were preloaded with the pH-sensitive LysoSensor Yellow/Blue Dextran 10kD (Invitrogen) based on the method described by Jonscher et al. (Jonscher et al., 2019). Briefly, fresh RBCs (stored less than a week) were washed three times in cold PBS. Of this pellet, 64 μL packed RBCs were transferred to 128 μL of freshly prepared preloading lysis buffer (5 mM K_2_HPO_4_, 20 mM Glucose, pH 7.4) and 2 μL 30mM DTT, 4 μL 50mM Mg-ATP (Sigma) and 2 μL of 50mg/mL LysoSensor Blue/Yellow dextran 10kD (Invitrogen) on ice. The suspension was rotated at 4°C for 10min. For resealing, 50 μL of 5× resealing buffer (750 mM NaCl, 25 mM Na_2_HPO_4_, pH 7.4) was carefully added to the RBC suspension and incubated for 60 min at 37°C while gently rocking. Preloaded RBCs were washed three times in cRPMI and resuspended in 1 mL cRPMI.

For infection of preloaded RBCs, schizonts were isolated from 30 mL of a 4% hematocrit highly synchronous, late stage *P. falciparum* 3D7 culture using a Percoll gradient. The resulting schizont pellet was resuspended in 1 mL cRPMI, mixed with the 1 mL preloaded RBCs and cultured for 20h at 37°C, 5% O_2_, 5% CO_2_ to allow re-invasion and formation of rings. Successful re-invasion was controlled with Giemsa-stained blood smears and flow cytometric measurement as described above. At ring stage, the culture was split into two parts and treated with either 2 μM Darapladib or DMSO control and further incubated until control parasites progressed to late trophozoite stage (32-40hpi, large hemozoin crystals visible). To identify infected RBCs, the cells were stained with 0.5 μM SYTO 61 (Invitrogen) for 30min at 37°C and washed once in cRPMI. The resulting pellet was resuspended in cRPMI and transferred onto a glass slide and covered with a glass cover slip. Images were taking with Deltavision Elite high resolution fluorescence microscope (GE Healthcare Lifesciences): LysoSensor pH 7, λ_Ex_ 390/18 nm, λ_Em_ 435/48, “Blue”; LysoSensor pH 5, λ_Ex_ 390/18 nm, λ_Em_ 597/45, “Yellow”; SYTO 61, λ_Ex_ 632/22 nm, λ_Em_ 679/34 and differential interference contrast (DIC). Fluorescent signal in the FV was quantified as the integrated density of the FV area in the “Yellow” and “Blue” channels using ImageJ. Relative successful transport of cytosolic LysoSensor to the FV was expressed as the ratio of integrated densities of “Yellow” and “Blue” channels.

For flow cytometric measurement, the cells were stained with 0.5 μM SYTO 61 for 30 min at 37°C, washed once in cRPMI afterwards and measured on a MACSquant flow cytometer: LysoSensor pH 7, λ_Ex_ 405 nm, λ_Em_ 450/50 nm, “Blue”; LysoSensor pH 5, λ_Ex_ 405 nm, λ_Em_ 525/50 nm, “Yellow” and SYTO 61, λ_Ex_ 561 nm, λ_Em_ 695/50 nm. Data was analyzed using FlowJo 10 (FlowJo, LLC). Briefly, after exclusion of doublets, debris and merozoites by size, infected RBCs were gated for by selecting SYTO 61 positive cells. The MFI of “Yellow” and “Blue” channels of SYTO 61^+^infected RBCs was calculated. Relative successful transport of cytosolic LysoSensor to the FV was expressed as the ratio of the MFI of “Yellow” and “Blue” channels.

### Artemisinin ring stage survival assay

Ring stage survival upon treatment with Dihydroartemisinin (DHA, Cayman Chemical) was measured based on the method described by Witkowski et al. (Witkowski et al., 2013). Briefly, ring stage survival was assessed in *P. falciparum* NF54 K13 C580Y and ART-resistant *P. falciparum* field isolate strains 3815 and 3801. Late stage schizonts were purified from synchronous *P. falciparum* cultures which contained only late stage (44–48 hpi) schizonts and newly invaded rings. Purified schizonts were added to fresh RBCs and the culture was incubated for 3 h at 37°C, to allow egress, re-invasion and development of rings. After 3h, unegressed schizonts were removed by a second Percoll purification. The pellet containing highly synchronous 0–3 h old rings and RBCs was washed in cRPMI and was examined by flow cytometry and Giemsa-stained blood smears to confirm schizont depletion and to measure the parasitemia. Subsequently, the parasitemia was adjusted to 1% and the culture diluted 1:10 in cRPMI. Initial parasitemia was measured by flow cytometry as described above and Giemsa-stained blood smears. In a 24-well plate, 1 mL aliquots of the cultures were treated with 700 nM DHA, 1μM Darapladib, or combinations of both drugs for 6h at 37°C, 5% O_2_, 5% CO_2_. The concentration of DMSO was kept equal in all samples. After 6 h, the drugs were removed by washing twice in cRPMI. After resuspension in cRPMI, the cultures were incubated for further 66 h and parasitemia was assessed by flow cytometry using SYBR Green I as described before and Giemsa-stained blood smears. Parasite survival was calculated as the ratio of rings in the treated and control samples after subtraction of the initial amount of rings (T-I/C-I).

## QUANTIFICATION AND STATISTICAL ANALYSIS

Statistical details are given in figure legends. If not otherwise indicated, at least three independent experiments (biological replicates) were performed. Statistical significance was determined using unpaired t-test. P values > 0.05 were considered as not significant. Data is presented as means ± SD. Statistical analysis was performed using GraphPad Prism (v9.2.0).

