## Supplemental Figures and Tables for "Human peroxiredoxin 6 is essential for malaria parasites and provides a host-based drug target"

### for

**Content:**

Supplementary Figures
Supplementary Tables

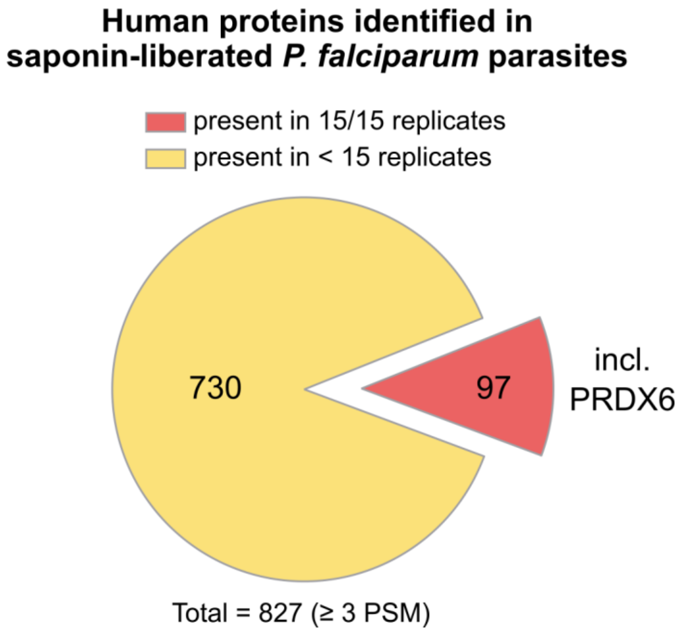


**Fig. S1. PRDX6 is identified in lysates of *P. falciparum* blood stage parasites.** *P. falciparum* infected RBCs were treated with saponin to selectively lyse the RBC membrane. The RBC cytosol was removed by centrifugation. In 15 biological replicates of LC-MS/MS analyses of saponin-liberated mature parasite lysates, 827 human proteins with ≥ 3 peptide spectrum matches (PSM) were detected and were subsequently filtered based on the frequency of occurrence across the 15 sample preparations. Only 97 of these human proteins, including PRDX6, were found in all 15 datasets. The full list of the 97 proteins with abundance information can be found in supplementary Data S1.

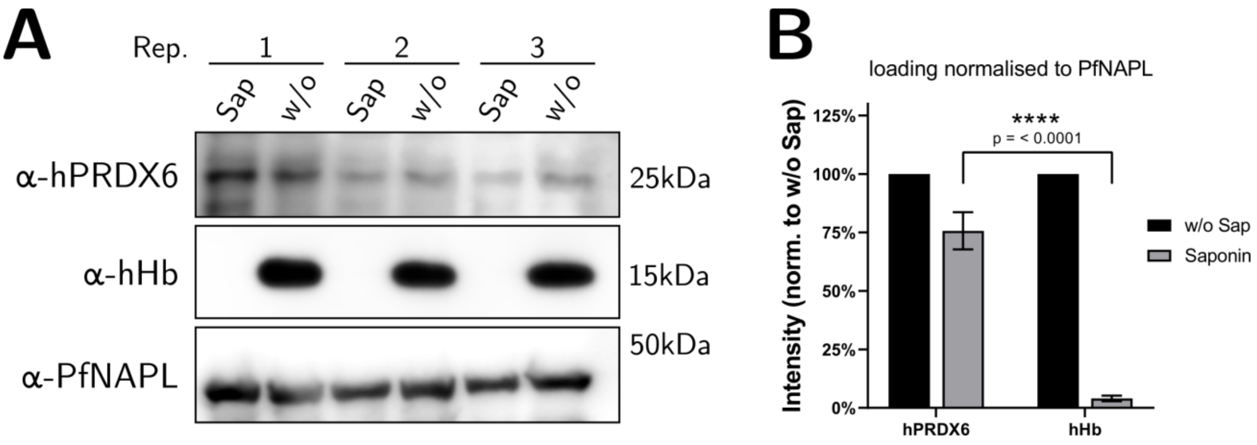


**Fig. S2. Host PRDX6 is internalized by *P. falciparum* blood stage parasites.** Immunoblots of saponin-treated and untreated schizonts show localization of human PRDX6 within the parasite. Saponization causes the loss of RBC cytosol proteins such as hemoglobin (Hb). *Pf*NAPL serves as a loading control. Band intensities for human PRDX6 and hemoglobin were quantified with the ImageJ software and normalized to *Pf*NAPL. Abb.: Sap, saponin; w/o, without saponin

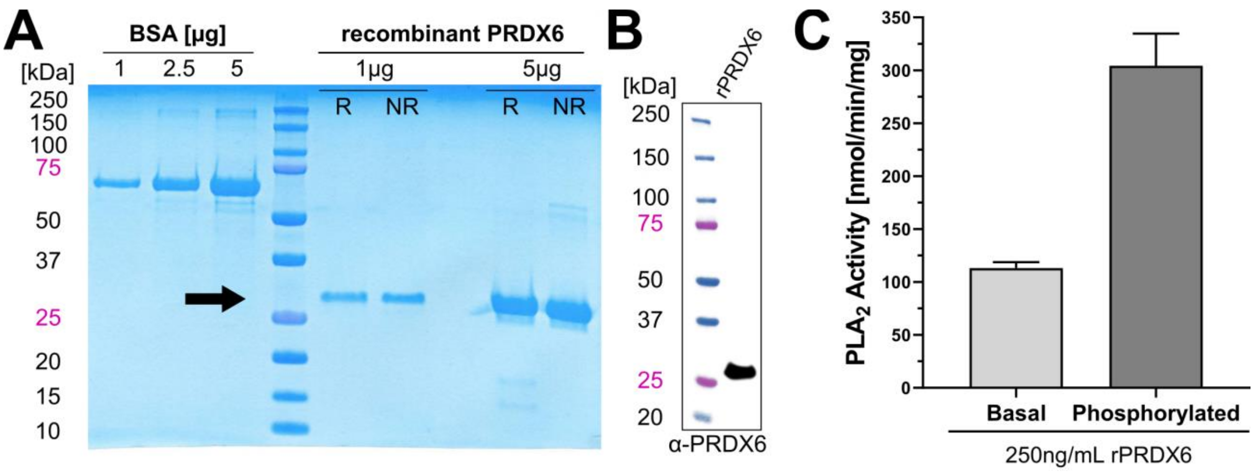


**Fig. S3. Expression and characterization of recombinant PRDX6 (rPRDX6).** (**A**) Purified rPRDX6 migrates as a single band at the expected size of 27 kDa on SDS-PAGE. (**B**) Identity of rPRDX6 was confirmed by immunoblotting with a monoclonal α-hPRDX6 antibody. (**C**) Specific $PLA_2$ activity of basal and phosphorylated rPRDX6 was consistent with published data (Wu et al., 2009).

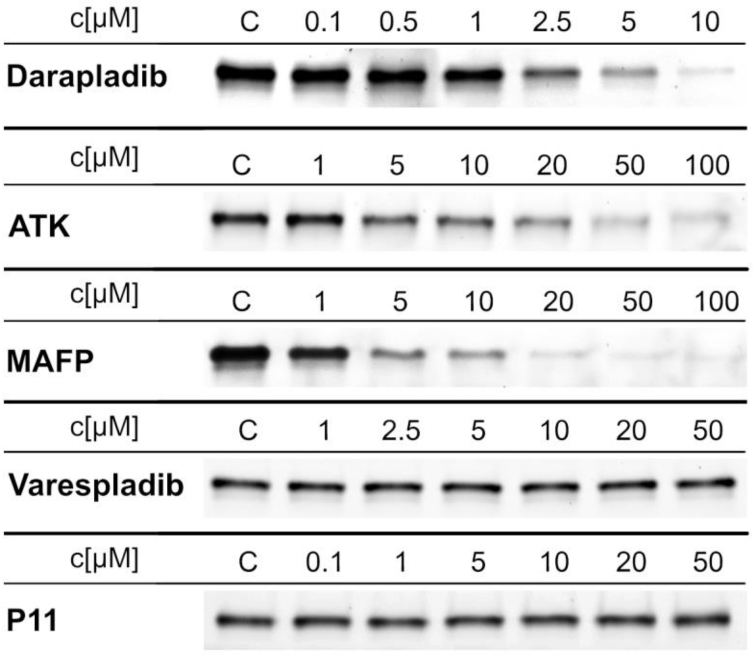


**Fig. S4. Darapladib, MAFP and ATK, but not Varespladib bind to human PRDX6.** Recombinant human PRDX6 was incubated with $PLA_2$ inhibitors Darapladib, ATK, MAFP, P11 and Varespladib and subsequently labelled with the fluorescent ABPP-probe TAMRA-FP. After SDS-PAGE, the polyacrylamide gel was imaged using a Cy3 filter set. Darapladib, MAFP and ATK, but not Varespladib and P11, reduced labelling of human PRDX6. This indicates that Darapladib, ATK and MAFP bind to the $PLA_2$ active site of PRDX6. Representative gels of three independent experiments are shown. Results from densitometry analysis of three independent experiments is shown in Figure 2A.

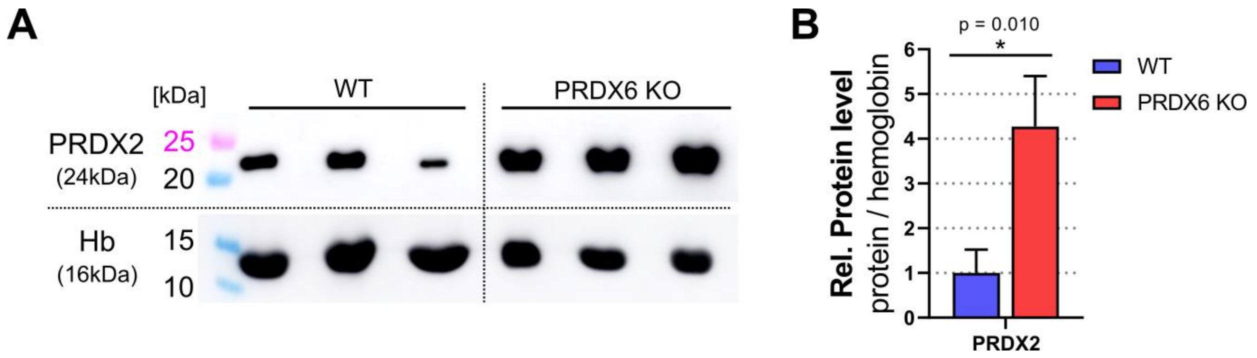


**Fig. S5. Loss of PRDX6 induces compensatory overexpression of PRDX2 in mice.** (**A**) The protein levels of PRDX2 and hemoglobin (control) were determined by immunoblotting of RBCs from *prdx6*$^{-/-}$ and WT mice. (**B**) Bands for PRDX2 from the immunoblot were quantified and normalized to hemoglobin levels by ImageJ software. Data are means ± SD of three independent experiments, unpaired t-test.

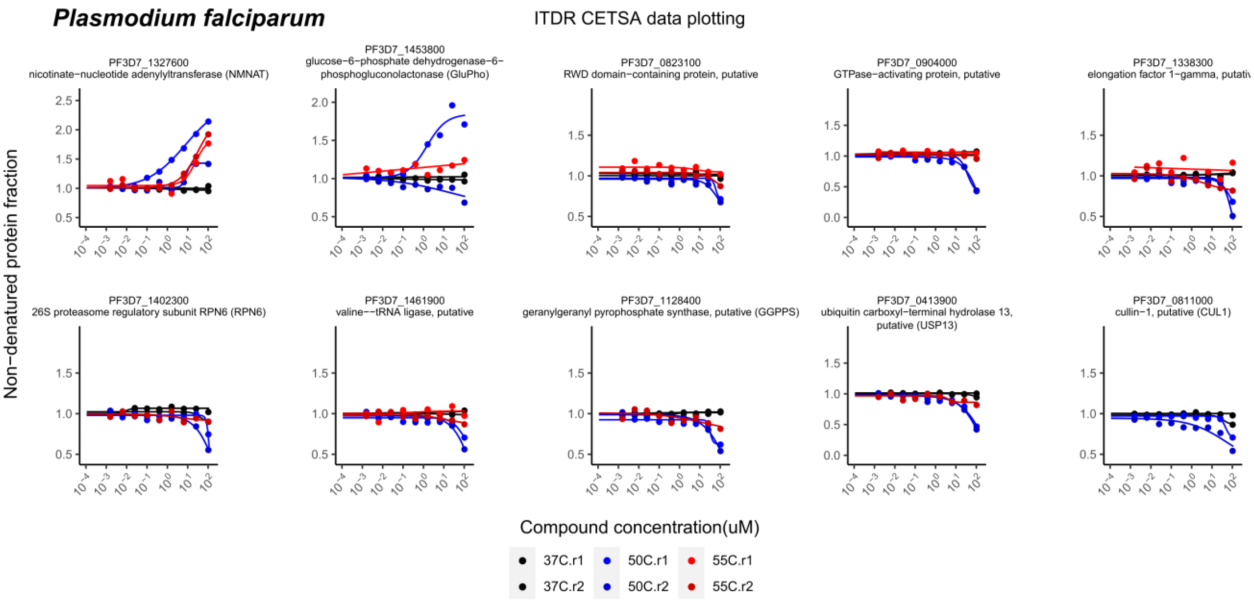


**Fig. S6. Thermal stability profile of proteins identified in ITDR-CETSA under Darapladib treatment.** Individual stability profiles of hits from Fig. 2G for the *P. falciparum* proteome. Soluble protein abundance after thermal challenge [50°C (blue) and 55°C (red)] is plotted relative to no-drug control along drug concentration gradient (x-axis). Non-denaturing control (37°C) plotted in black. Data represents 2 independent biological replicates.

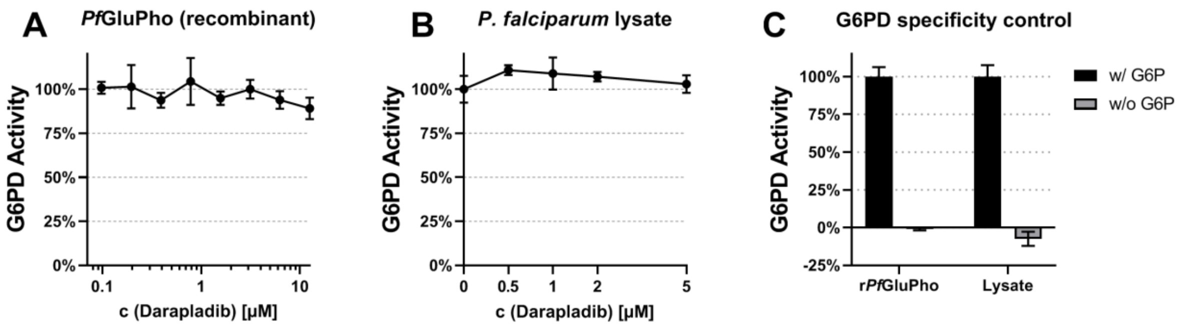


**Fig. S7. Darapladib does not inhibit the essential glucose-6-phosphate dehydrogenase (G6PD) activity of *Pf*GluPho. (A)** The glucose-6-phosphate dehydrogenase (G6PD) activity of recombinant *P. falciparum* glucose-6-phosphate dehydrogenase-6-phosphogluconolactonase (*Pf*GluPho) was measured by quantification of NADPH production. Darapladib does not inhibit the G6PD activity of recombinant *Pf*GluPho. (**B**) Darapladib does not block G6PD activity in *P. falciparum* 3D7 lysates. (**C**) No G6PD background activity or non-specific NADPH production was detected in control samples which lacked the substrate glucose-6-phosphate. Data are means ± SD of two independent experiments.

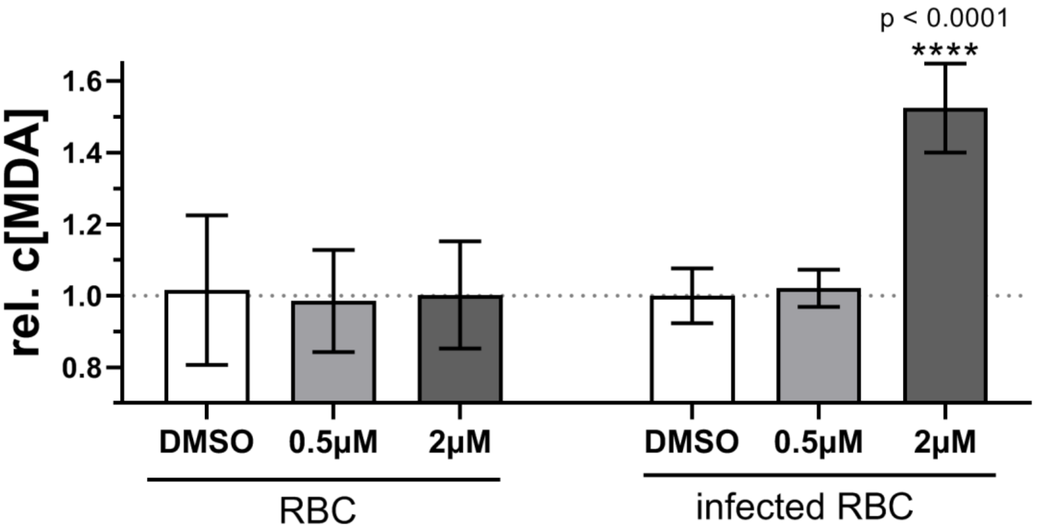


**Fig. S8. Darapladib treatment increases malondialdehyde (MDA) levels.** Treatment of asynchronous *P. falciparum* 3D7 cultures with Darapladib lead to a significant increase of MDA levels in infected RBCs as detected with the TBARS assay. Uninfected RBCs remained unaffected following Darapladib treatment. Data are means ± SD of three independent experiments, unpaired t-test.

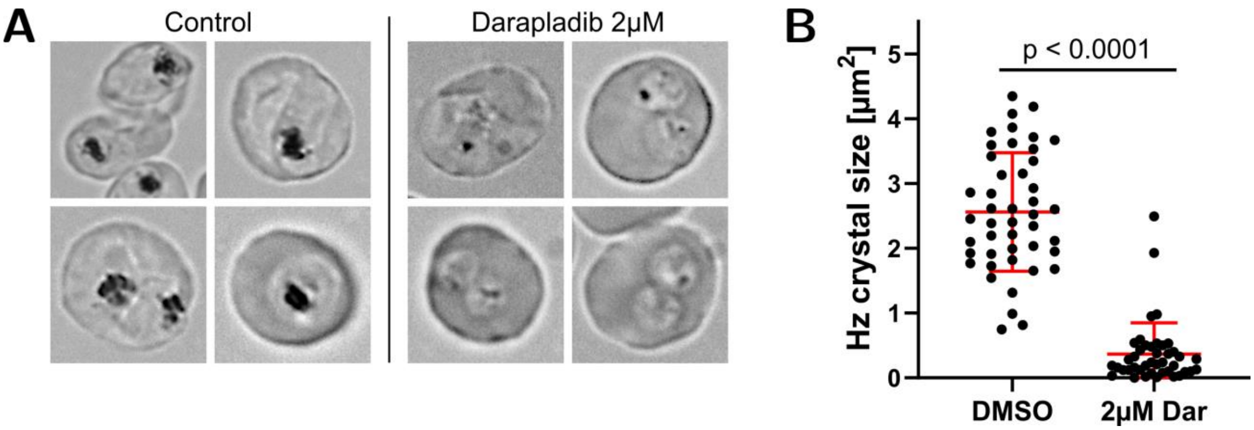


**Fig. S9. Treatment of *P. falciparum* with Darapladib at ring stage decreases hemozoin crystal size.** (**A**) DIC images show that *P. falciparum* 3D7 parasites contain smaller hemozoin crystals following treatment with Darapladib. Representative images from three independent experiments are shown. (**B**) The average size of hemozoin crystals was measured using ImageJ and is greatly reduced by treatment with Darapladib. Data are means ± SD of three independent experiments, unpaired t-test.

#### SUPPLEMENTARY TABLES

##### Table S1.

**$IC_{50}$ values for several $PLA_2$ inhibitors for inhibition of *P. falciparum* blood stage progression and growth.** Data corresponds to Figure 2B in the main text. Synchronous *P. falciparum* 3D7 culture were treated at ring stage with inhibitors and subsequently allowed to progress to schizont stage ("progression") or to complete the full blood stage cycle until the next generation ring stage ("growth") and measured quantitatively by flow cytometry using SYBR Green I. Data are means ± SD of three independent experiments. ***Abb.:*** *ATK, Arachidonyl trifluoromethyl ketone; MAFP, Methoxy arachidonyl fluorophosphonate*

| Inhibitor | $IC_{50}$ Progression [µM] | $IC_{50}$ Growth [µM] |
|---|---|---|
| Darapladib | 0.56 ± 0.02 | 0.76 ± 0.01 |
| ATK | 3.63 ± 0.1 | 2.14 ± 0.06 |
| MAFP | 9.09 ± 0.18 | 3.86 ± 0.08 |
| Varespladib | No inhibition | No inhibition |
| P11 | No inhibition | No inhibition |

##### Table S2.

**Co-treatment with Darapladib and Artemisinin synergistically reduces parasite survival.** The data corresponds to Figure 4A in the main text. The survival rate for treatment with 1 µM Darapladib alone and 700 nM DHA alone was multiplied to calculate a theoretical survival rate if the effect of the combination was additive ("Calc. Additive"). The experimentally measured survival rate ("Measured") was divided by the calculated theoretical additive survival rate for the co-treatment. A ratio < 1 for Calc./Measured indicates synergism, a ratio = 1 indicates an additive effect and a ratio > 1 indicates an antagonistic effect. Data are means ± SD of three independent experiments, unpaired t-test.

| | **Parasite survival** | | | | |
|---|---|---|---|---|---|
| | **Drugs Alone** | | **Combination** | | **Ratio** |
| **Strain** | **Dar 1µM** | **DHA** | **Calc. Additive** | **Measured** | **Measur./Calc.** |
| NF54 C580Y | 82.3 ± 5.3% | 7.1 ± 1.8% | 5.8 ± 1.9% | 0.5 ± 2.2% | 0.09 * (p=0.03) |
| 3815 | 94.7 ± 3.2% | 7.1 ± 0.9% | 6.7 ± 1.1% | 3.3 ± 0.5% | 0.49 ** (p=0.008) |
| 3601 | 89.6 ± 6.7% | 28.3 ± 3.0% | 25.4 ± 4.6% | 21.9 ± 2.7% | 0.89 ns |

#### SUPPLEMENTAL ITEM TITLE AND LEGEND

##### Data S1.

Supplementary DATA S1 MS internalized human proteins.pdf

**PRDX6 is identified in lysates of blood stage *P. falciparum* parasites.** *P. falciparum* infected RBCs were treated with saponin to selectively lyse the RBC membrane. 4242 human proteins were detected in LC-MS/MS analyses of saponin-liberated mature parasite lysates in 15 biological replicates. 827 of the human proteins, which were found with ≥3 peptide spectrum matches (PSM), were subsequently filtered based on the frequency of occurrence across the 15 sample preparations. Only 97 of these human proteins, including PRDX6, were found in all 15 datasets.
