## Supplemental Data S1 for "Human peroxiredoxin 6 is essential for malaria parasites and provides a host-based drug target"

**Legend**: 97 human proteins consistently identified (PSM≥3) in LC-MS/MS analysis of Saponin-liberated mature P.falciparum trophozoite lysates (15/15)

| ID | Protein Name | Rep1 #PSM | Rep2 #PSM | Rep3 #PSM | Rep4 #PSM | Rep5 #PSM | Rep6 #PSM | Rep7 #PSM | Rep8 #PSM | Rep9 #PSM | Rep10 #PSM | Rep11 #PSM | Rep12 #PSM | Rep13 #PSM | Rep14 #PSM | Rep15 #PSM | #PSM Average | #PSM stdev |
|---|---|---|---|---|---|---|---|---|---|---|---|---|---|---|---|---|---|---|
| P68871 | Hemoglobin subunit beta | 1509 | 1868 | 1552 | 1457 | 1407 | 1339 | 797 | 449 | 394 | 699 | 539 | 414 | 547 | 603 | 760 | 956 | 504 |
| P69905 | Hemoglobin subunit alpha | 1638 | 1347 | 1327 | 1118 | 1484 | 1384 | 430 | 284 | 306 | 218 | 416 | 175 | 218 | 317 | 484 | 743 | 557 |
| P02042 | Hemoglobin subunit delta | 901 | 981 | 918 | 884 | 833 | 819 | 479 | 293 | 259 | 367 | 320 | 216 | 347 | 368 | 501 | 566 | 285 |
| P02549 | Spectrin alpha chain, erythrocytic 1 | 352 | 294 | 306 | 260 | 306 | 319 | 292 | 407 | 384 | 159 | 222 | 146 | 99 | 227 | 292 | 271 | 87 |
| P00915 | Carbonic anhydrase 1 | 374 | 366 | 350 | 341 | 350 | 314 | 262 | 150 | 136 | 140 | 219 | 124 | 349 | 233 | 314 | 268 | 94 |
| P04406 | Glyceraldehyde-3-phosphate dehydrogenase | 360 | 311 | 301 | 269 | 296 | 280 | 297 | 303 | 316 | 122 | 331 | 179 | 158 | 166 | 209 | 260 | 73 |
| P04040 | Catalase | 236 | 207 | 193 | 228 | 192 | 192 | 154 | 155 | 149 | 116 | 156 | 118 | 163 | 231 | 257 | 183 | 43 |
| P11142 | Heat shock cognate 71 kDa protein | 162 | 194 | 163 | 142 | 164 | 139 | 176 | 238 | 222 | 101 | 155 | 131 | 139 | 174 | 159 | 164 | 35 |
| Q13228-4 | Isoform 4 of Methanethiol oxidase | 140 | 113 | 117 | 128 | 130 | 130 | 122 | 174 | 160 | 63 | 123 | 78 | 84 | 104 | 123 | 119 | 29 |
| P69891 | Hemoglobin subunit gamma-1 | 219 | 208 | 221 | 200 | 209 | 189 | 42 | 54 | 49 | 46 | 73 | 46 | 43 | 52 | 80 | 115 | 79 |
| P11171 | Protein 4.1 | 133 | 120 | 121 | 116 | 131 | 121 | 124 | 116 | 106 | 82 | 113 | 76 | 107 | 100 | 135 | 113 | 17 |
| P62979 | Ubiquitin-40S ribosomal protein S27a | 122 | 103 | 107 | 117 | 110 | 98 | 99 | 130 | 119 | 96 | 127 | 97 | 105 | 134 | 115 | 112 | 13 |
| P04264 | Keratin, type II cytoskeletal 1 | 96 | 126 | 98 | 141 | 86 | 156 | 99 | 89 | 115 | 50 | 122 | 78 | 37 | 88 | 69 | 97 | 32 |
| P60709 | Actin, cytoplasmic 1 | 106 | 92 | 79 | 72 | 86 | 84 | 92 | 133 | 115 | 52 | 119 | 70 | 70 | 123 | 149 | 96 | 27 |
| P32119 | Peroxiredoxin-2 | 90 | 110 | 105 | 104 | 93 | 92 | 61 | 62 | 54 | 75 | 77 | 62 | 148 | 123 | 175 | 95 | 34 |
| P00441 | Superoxide dismutase [Cu-Zn] | 123 | 115 | 108 | 112 | 100 | 104 | 58 | 101 | 83 | 58 | 73 | 77 | 107 | 78 | 92 | 93 | 20 |
| P35612 | Beta-adducin | 100 | 76 | 89 | 81 | 93 | 92 | 107 | 121 | 129 | 64 | 81 | 61 | 84 | 96 | 100 | 92 | 19 |
| P0DMV9 | Heat shock 70 kDa protein 1B | 98 | 102 | 84 | 84 | 90 | 89 | 91 | 105 | 97 | 71 | 77 | 80 | 78 | 119 | 95 | 91 | 13 |
| P13645 | Keratin, type I cytoskeletal 10 | 77 | 96 | 90 | 154 | 68 | 142 | 107 | 81 | 74 | 55 | 99 | 86 | 47 | 90 | 66 | 89 | 29 |
| P30043 | Flavin reductase (NADPH) | 111 | 98 | 95 | 71 | 102 | 88 | 47 | 60 | 66 | 51 | 76 | 31 | 78 | 55 | 88 | 74 | 23 |
| P60174 | Triosephosphate isomerase | 74 | 87 | 77 | 78 | 75 | 66 | 70 | 86 | 79 | 48 | 69 | 56 | 63 | 71 | 88 | 72 | 11 |
| P07384 | Calpain-1 catalytic subunit | 73 | 63 | 64 | 57 | 68 | 62 | 81 | 96 | 92 | 51 | 46 | 21 | 81 | 57 | 67 | 65 | 19 |
| P35908 | Keratin, type II cytoskeletal 2 epidermal | 64 | 66 | 54 | 103 | 58 | 114 | 84 | 48 | 39 | 51 | 82 | 58 | 26 | 66 | 47 | 64 | 23 |
| P07900-2 | Isoform 2 of Heat shock protein HSP 90-alpha | 58 | 54 | 63 | 55 | 61 | 55 | 65 | 63 | 57 | 44 | 51 | 56 | 70 | 77 | 87 | 61 | 11 |
| P35527 | Keratin, type I cytoskeletal 9 | 49 | 69 | 66 | 71 | 49 | 70 | 55 | 75 | 111 | 32 | 76 | 72 | 21 | 50 | 46 | 61 | 21 |
| P13798 | Acylamino-acid-releasing enzyme | 80 | 64 | 63 | 65 | 71 | 65 | 54 | 72 | 79 | 33 | 63 | 37 | 47 | 42 | 67 | 60 | 15 |
| P02730 | Band 3 anion transport protein | 44 | 40 | 35 | 34 | 52 | 47 | 52 | 120 | 125 | 26 | 71 | 54 | 33 | 48 | 107 | 59 | 32 |
| P06753-2 | Isoform 2 of Tropomyosin alpha-3 chain | 58 | 58 | 57 | 78 | 61 | 55 | 51 | 78 | 72 | 21 | 48 | 38 | 27 | 54 | 60 | 54 | 16 |
| P00918 | Carbonic anhydrase 2 | 94 | 87 | 90 | 83 | 93 | 76 | 73 | 21 | 24 | 25 | 25 | 18 | 8 | 27 | 38 | 52 | 33 |
| P62258 | 14-3-3 protein epsilon | 62 | 66 | 58 | 52 | 49 | 51 | 52 | 69 | 55 | 42 | 44 | 41 | 34 | 48 | 56 | 52 | 10 |
| P02100 | Hemoglobin subunit epsilon | 94 | 78 | 95 | 78 | 99 | 97 | 18 | 32 | 26 | 19 | 40 | 23 | 23 | 23 | 24 | 51 | 34 |
| P02768 | Serum albumin | 41 | 63 | 43 | 30 | 31 | 236 | 34 | 18 | 36 | 31 | 61 | 42 | 11 | 43 | 47 | 51 | 53 |
| P06733 | Alpha-enolase | 73 | 48 | 54 | 49 | 59 | 51 | 60 | 39 | 32 | 18 | 53 | 35 | 61 | 57 | 57 | 50 | 14 |
| P63104 | 14-3-3 protein zeta/delta | 55 | 59 | 43 | 46 | 43 | 41 | 42 | 70 | 52 | 42 | 52 | 47 | 36 | 57 | 58 | 50 | 9 |
| P00390 | Glutathione reductase, mitochondrial | 50 | 51 | 47 | 37 | 47 | 50 | 44 | 62 | 60 | 20 | 54 | 42 | 34 | 48 | 39 | 46 | 11 |
| P31946 | 14-3-3 protein beta/alpha | 52 | 57 | 45 | 42 | 41 | 38 | 35 | 67 | 47 | 36 | 43 | 39 | 33 | 48 | 47 | 45 | 9 |
| P14625 | Endoplasmin | 40 | 40 | 38 | 36 | 40 | 34 | 37 | 66 | 52 | 47 | 50 | 42 | 51 | 45 | 51 | 45 | 8 |
| Q08495 | Dematin | 40 | 45 | 38 | 37 | 47 | 37 | 39 | 49 | 48 | 40 | 41 | 31 | 59 | 45 | 55 | 43 | 7 |
| P61981 | 14-3-3 protein gamma | 44 | 57 | 41 | 39 | 38 | 36 | 33 | 64 | 47 | 40 | 40 | 34 | 33 | 42 | 43 | 42 | 9 |
| P55072 | Transitional endoplasmic reticulum ATPase | 63 | 52 | 51 | 53 | 47 | 52 | 55 | 34 | 25 | 20 | 32 | 17 | 20 | 39 | 45 | 40 | 15 |
| Q06830 | Peroxiredoxin-1 | 51 | 41 | 54 | 52 | 38 | 39 | 28 | 28 | 31 | 31 | 30 | 29 | 32 | 48 | 54 | 39 | 10 |
| P40925-3 | Isoform 3 of Malate dehydrogenase, cytoplasmic | 47 | 44 | 38 | 36 | 37 | 33 | 31 | 38 | 39 | 32 | 36 | 37 | 34 | 45 | 45 | 38 | 5 |
| P07195 | L-lactate dehydrogenase B chain | 63 | 62 | 49 | 48 | 48 | 44 | 39 | 22 | 22 | 11 | 27 | 21 | 21 | 35 | 40 | 37 | 16 |
| P23526 | Adenosylhomocysteinase | 44 | 40 | 40 | 44 | 46 | 47 | 35 | 30 | 34 | 20 | 37 | 23 | 19 | 40 | 42 | 36 | 9 |
| P55786 | Puromycin-sensitive aminopeptidase | 45 | 33 | 37 | 35 | 42 | 35 | 26 | 40 | 52 | 20 | 42 | 24 | 21 | 26 | 36 | 34 | 9 |
| P00558 | Phosphoglycerate kinase 1 | 52 | 51 | 50 | 50 | 54 | 50 | 34 | 17 | 20 | 16 | 23 | 20 | 4 | 28 | 26 | 33 | 17 |
| P37837 | Transaldolase | 41 | 36 | 31 | 46 | 41 | 34 | 36 | 30 | 32 | 15 | 34 | 25 | 21 | 30 | 38 | 33 | 8 |
| P61970 | Nuclear transport factor 2 | 45 | 37 | 35 | 30 | 42 | 35 | 47 | 40 | 46 | 7 | 42 | 14 | 9 | 24 | 35 | 33 | 13 |
| P29144 | Tripeptidyl-peptidase 2 | 51 | 52 | 35 | 30 | 52 | 38 | 46 | 30 | 39 | 10 | 25 | 16 | 7 | 21 | 19 | 31 | 15 |
| P02533 | Keratin, type I cytoskeletal 14 | 29 | 36 | 33 | 42 | 24 | 57 | 31 | 23 | 45 | 26 | 32 | 35 | 10 | 23 | 15 | 31 | 12 |

| | | | | | | | | | | | | | | | | | | |
|---|---|---|---|---|---|---|---|---|---|---|---|---|---|---|---|---|---|---|
| P09104 | Gamma-enolase | 38 | 25 | 33 | 27 | 33 | 23 | 34 | 27 | 18 | 8 | 35 | 24 | 54 | 30 | 34 | 30 | 10 |
| P09493-5 | Isoform 5 of Tropomyosin alpha-1 chain | 35 | 37 | 31 | 39 | 31 | 31 | 33 | 32 | 33 | 11 | 26 | 21 | 10 | 31 | 35 | 29 | 9 |
| P07203 | Glutathione peroxidase 1 | 34 | 25 | 30 | 20 | 29 | 22 | 36 | 20 | 20 | 14 | 55 | 33 | 18 | 31 | 31 | 28 | 10 |
| P50395 | Rab GDP dissociation inhibitor beta | 39 | 44 | 40 | 38 | 37 | 33 | 21 | 14 | 16 | 16 | 26 | 14 | 11 | 27 | 40 | 28 | 12 |
| P35579 | Myosin-9 | 37 | 40 | 28 | 31 | 34 | 31 | 42 | 17 | 21 | 19 | 7 | 9 | 4 | 46 | 50 | 28 | 14 |
| Q9NY33 | Dipeptidyl peptidase 3 | 39 | 29 | 30 | 27 | 27 | 29 | 30 | 31 | 26 | 11 | 29 | 17 | 21 | 30 | 28 | 27 | 7 |
| P00491 | Purine nucleoside phosphorylase | 42 | 43 | 37 | 37 | 51 | 47 | 21 | 7 | 12 | 4 | 18 | 6 | 12 | 22 | 32 | 26 | 16 |
| P09211 | Glutathione S-transferase P | 29 | 27 | 22 | 24 | 31 | 24 | 16 | 44 | 43 | 6 | 26 | 17 | 29 | 20 | 33 | 26 | 10 |
| Q16775 | Hydroxyacylglutathione hydrolase, mitochondrial | 30 | 26 | 34 | 23 | 18 | 35 | 21 | 29 | 26 | 13 | 31 | 18 | 21 | 32 | 27 | 26 | 6 |
| Q13867 | Bleomycin hydrolase | 28 | 27 | 26 | 28 | 28 | 26 | 22 | 32 | 39 | 9 | 37 | 20 | 13 | 17 | 13 | 24 | 9 |
| P36959 | GMP reductase 1 | 25 | 23 | 29 | 21 | 27 | 28 | 22 | 24 | 29 | 14 | 24 | 12 | 23 | 21 | 36 | 24 | 6 |
| P07738 | Bisphosphoglycerate mutase | 45 | 29 | 35 | 34 | 41 | 36 | 29 | 7 | 10 | 9 | 11 | 6 | 7 | 20 | 26 | 23 | 14 |
| P62820 | Ras-related protein Rab-1A | 26 | 25 | 26 | 23 | 23 | 22 | 17 | 29 | 29 | 11 | 18 | 18 | 16 | 27 | 25 | 22 | 5 |
| P00492 | Hypoxanthine-guanine phosphoribosyltransferase | 21 | 20 | 28 | 20 | 26 | 25 | 23 | 17 | 16 | 17 | 22 | 12 | 18 | 23 | 32 | 21 | 5 |
| P04632 | Calpain small subunit 1 | 27 | 26 | 19 | 19 | 19 | 25 | 28 | 27 | 25 | 20 | 16 | 6 | 20 | 17 | 24 | 21 | 6 |
| Q00013 | 55 kDa erythrocyte membrane protein | 30 | 25 | 23 | 16 | 16 | 18 | 20 | 22 | 27 | 16 | 26 | 16 | 14 | 18 | 21 | 21 | 5 |
| Q99497 | Protein/nucleic acid deglycase DJ-1 | 36 | 40 | 32 | 28 | 30 | 32 | 20 | 6 | 5 | 11 | 10 | 7 | 10 | 20 | 20 | 20 | 12 |
| P30041 | Peroxiredoxin-6 | 28 | 26 | 26 | 26 | 24 | 28 | 21 | 6 | 8 | 16 | 10 | 8 | 13 | 22 | 26 | 19 | 8 |
| P62826 | GTP-binding nuclear protein Ran | 27 | 21 | 24 | 23 | 21 | 27 | 20 | 12 | 18 | 9 | 12 | 14 | 11 | 14 | 33 | 19 | 7 |
| P28289 | Tropomodulin-1 | 29 | 21 | 16 | 19 | 16 | 15 | 21 | 12 | 15 | 17 | 14 | 15 | 10 | 30 | 34 | 19 | 7 |
| P48637 | Glutathione synthetase | 20 | 22 | 19 | 18 | 22 | 19 | 17 | 20 | 18 | 7 | 20 | 12 | 16 | 23 | 29 | 19 | 5 |
| P60900 | Proteasome subunit alpha type-6 | 20 | 17 | 22 | 23 | 24 | 21 | 23 | 17 | 22 | 10 | 15 | 12 | 8 | 24 | 19 | 18 | 5 |
| P25786-2 | Isoform Long of Proteasome subunit alpha type-1 | 26 | 21 | 24 | 19 | 24 | 23 | 20 | 11 | 12 | 7 | 12 | 10 | 7 | 16 | 26 | 17 | 7 |
| P25788 | Proteasome subunit alpha type-3 | 19 | 17 | 16 | 17 | 19 | 22 | 19 | 16 | 17 | 11 | 19 | 12 | 8 | 21 | 22 | 17 | 4 |
| P28066 | Proteasome subunit alpha type-5 | 23 | 19 | 19 | 13 | 30 | 22 | 19 | 18 | 15 | 10 | 13 | 7 | 9 | 14 | 16 | 16 | 6 |
| P07477 | Trypsin-1 | 15 | 13 | 16 | 17 | 16 | 8 | 13 | 15 | 23 | 15 | 25 | 15 | 12 | 27 | 15 | 16 | 5 |
| Q16881 | Thioredoxin reductase 1, cytoplasmic | 21 | 20 | 19 | 22 | 17 | 18 | 14 | 19 | 19 | 8 | 15 | 16 | 6 | 15 | 14 | 16 | 4 |
| O14818 | Proteasome subunit alpha type-7 | 17 | 14 | 17 | 19 | 20 | 18 | 15 | 11 | 13 | 10 | 17 | 7 | 9 | 21 | 18 | 15 | 4 |
| Q9BRF8 | Serine/threonine-protein phosphatase CPPED1 | 19 | 15 | 15 | 15 | 17 | 15 | 18 | 18 | 20 | 5 | 12 | 8 | 8 | 17 | 15 | 14 | 4 |
| P17174 | Aspartate aminotransferase, cytoplasmic | 20 | 17 | 15 | 21 | 19 | 15 | 14 | 8 | 12 | 7 | 16 | 10 | 12 | 13 | 14 | 14 | 4 |
| P22392-2 | Isoform 3 of Nucleoside diphosphate kinase B | 22 | 19 | 19 | 20 | 22 | 18 | 15 | 7 | 7 | 6 | 7 | 4 | 7 | 9 | 18 | 13 | 7 |
| P28074 | Proteasome subunit beta type-5 | 19 | 15 | 18 | 13 | 16 | 13 | 15 | 8 | 14 | 6 | 15 | 9 | 7 | 15 | 12 | 13 | 4 |
| P11586 | C-1-tetrahydrofolate synthase, cytoplasmic | 14 | 18 | 14 | 14 | 13 | 11 | 13 | 15 | 18 | 9 | 8 | 10 | 5 | 13 | 14 | 13 | 4 |
| P20618 | Proteasome subunit beta type-1 | 16 | 11 | 11 | 19 | 14 | 11 | 12 | 14 | 16 | 6 | 9 | 12 | 7 | 16 | 12 | 12 | 4 |
| P00568 | Adenylate kinase isoenzyme 1 | 19 | 17 | 19 | 14 | 16 | 16 | 17 | 7 | 4 | 9 | 8 | 9 | 4 | 6 | 12 | 12 | 5 |
| P78417 | Glutathione S-transferase omega-1 | 20 | 16 | 20 | 15 | 14 | 18 | 12 | 7 | 6 | 6 | 5 | 11 | 7 | 7 | 12 | 12 | 5 |
| P62834 | Ras-related protein Rap-1A | 12 | 12 | 11 | 13 | 14 | 12 | 9 | 10 | 12 | 5 | 8 | 7 | 8 | 15 | 17 | 11 | 3 |
| P53004 | Biliverdin reductase A | 16 | 12 | 16 | 10 | 15 | 12 | 12 | 7 | 10 | 5 | 10 | 9 | 5 | 10 | 12 | 11 | 3 |
| P10599 | Thioredoxin | 7 | 13 | 7 | 9 | 13 | 9 | 6 | 11 | 11 | 8 | 10 | 7 | 26 | 8 | 12 | 10 | 5 |
| Q04760 | Lactoylglutathione lyase | 15 | 14 | 14 | 13 | 17 | 14 | 9 | 7 | 10 | 4 | 12 | 6 | 5 | 6 | 9 | 10 | 4 |
| P11166 | Solute carrier family 2, facilitated glucose transporter memb | 9 | 10 | 8 | 7 | 12 | 8 | 9 | 19 | 23 | 7 | 9 | 8 | 4 | 8 | 11 | 10 | 5 |
| P35754 | Glutaredoxin-1 | 10 | 10 | 7 | 11 | 9 | 8 | 10 | 11 | 10 | 4 | 6 | 4 | 11 | 11 | 25 | 10 | 5 |
| P28072 | Proteasome subunit beta type-6 | 12 | 13 | 10 | 13 | 13 | 8 | 6 | 8 | 10 | 7 | 4 | 11 | 7 | 11 | 11 | 10 | 3 |
| Q9UKK9 | ADP-sugar pyrophosphatase | 12 | 12 | 8 | 8 | 11 | 8 | 7 | 12 | 12 | 6 | 10 | 7 | 4 | 11 | 15 | 10 | 3 |
| Q13126-2 | Isoform 2 of S-methyl-5'-thioadenosine phosphorylase | 12 | 12 | 8 | 6 | 10 | 10 | 6 | 10 | 10 | 6 | 9 | 6 | 6 | 7 | 9 | 8 | 2 |
| P12955 | Xaa-Pro dipeptidase | 12 | 11 | 10 | 7 | 9 | 9 | 10 | 5 | 5 | 5 | 6 | 7 | 6 | 11 | 11 | 8 | 3 |
| P63208 | S-phase kinase-associated protein 1 | 12 | 8 | 10 | 7 | 6 | 11 | 10 | 7 | 7 | 4 | 6 | 5 | 5 | 9 | 9 | 8 | 2 |
